# A novel hypoxia gene signature indicates prognosis and reveals the multi-omics molecular landscape of tumour tissue in patients with hepatocellular carcinoma

**DOI:** 10.1101/2020.07.10.198176

**Authors:** Qiangnu Zhang, Juan Liao, Lijun Qiao, Quan Liu, Pengyu Liu, Mengting Xia, Xiaotao Huang, Liping Liu

**Affiliations:** Department of Hepatobiliary and Pancreas Surgery, Shenzhen People’s Hospital (The Second Clinical Medical College, Jinan University; The First Affiliated Hospital, Southern University of Science and Technology), Shenzhen 518020, Guangdong, China; Integrated Chinese and Western Medicine Postdoctoral Research Station, Jinan University, Guangzhou, Guangdong 510632, China; Department of Gastroenterology, West China School of Public Health and West China Fourth Hospital, Sichuan University, Chengdu610041, China; Department of Gastroenterology and Hepatology, Erasmus MC-University Medical Center, Rotterdam; Department of Clinical Medicine, North Sichuan Medical College, Nanchong637100, China

**Keywords:** HCC, hypoxia, gene signature, multi-omics, prognosis, microenvironment

## Abstract

**Background:** Previous studies on the impact of hypoxia on hepatocellular carcinoma (HCC) mostly focused on in vitro and animal models. The clinical value of assessing the degree of hypoxia in in vivo tissues and hypoxia-related molecular landscapes in HCC remain poorly defined.

**Methods:** A novel hypoxia gene signature was extracted from hypoxia-treated HCC cells using microarray analysis and a robust rank aggregation algorithm and was verified using public data. Next, the relationships between the hypoxia gene signature and the clinical characteristics and prognoses of HCC patients were analysed. Based on the multi-omics data from The Cancer Genome Atlas-Liver Hepatocellular Carcinoma (TCGA-LIHC) and 10 independent HCC cohorts from the Gene Expression Omnibus (GEO), a comprehensive analysis was conducted using the hypoxia gene signature to describe hypoxia-associated multi-omic molecular landscapes and immune microenvironments in HCC tissues.

**Results:** A 21-gene hypoxia signature was constructed to effectively indicate the exposure of hypoxia in HCC tissues. This novel hypoxia signature and the hypoxia scores calculated based on this signature were closely correlated to some clinical characteristics of HCC patients and can be used to evaluate prognosis. HCC tissues with different hypoxia scores differed significantly in transcriptomic, genomic, epigenomic, and proteomic alterations and immune microenvironments, some of which were related to the clinical prognoses of patients.

**Conclusion:** The 21-gene signature can effectively estimate hypoxia exposure of HCC tissues and has clinical value in the assessment and prediction of the prognosis of HCC patients. Using this 21-gene signature, hypoxia-associated molecular landscapes were described at the tissue level. This comprehensive molecular-level understanding can help to further elucidate the mechanism of hypoxia in tumours and guide clinical cancer therapy. The assessment of the degree of hypoxia is strongly recommended in the personalized treatment of HCC patients to benefit specific patient groups.

## Background

In 2018, there were 841,080 new cases of liver cancer worldwide, ranking sixth in the global incidence of all cancers. In the same year, liver cancer caused 781,631 deaths worldwide, ranking as the fourth leading cause of cancer death in the world [1]. Hepatocellular carcinoma (HCC) accounts for 85%-95% of primary liver cancers[2]. Due to the insidious onset and inadequate early diagnosis measures, 80% of patients are already in the middle and late stages of disease at the time of diagnosis, thus missing the optimal time for surgery[3]. In patients with advanced HCC, the mortality rate is as high as 80%, the median survival is less than 1 year, and the 5-year survival rate is less than 20% [4]. Although the development of targeted therapy and immunotherapy for the treatment of HCC has brought new hope to patients with advanced HCC, the overall efficacy of these therapeutic methods remains dismal[5]. The initiation and progression of HCC involves interactions with microenvironments. Therefore, revealing the diversity of characteristics of tumour cells brought by microenvironments will be conducive to the early diagnosis of HCC, to finding treatment targets, to establishing personalized treatment, and to predicting prognosis[6].

Hypoxia is the most important characteristic of solid tumour microenvironments. It leads to increased invasiveness, increased angiogenesis, increased stemness, metabolic reprogramming, changes in immune response, and resistance to radiochemotherapy [7, 8]. Because the assessment of hypoxia exposure in cancer tissues in vivo or in vitro is not very convenient, most studies on hypoxia in cancer are based on in vitro cell experiments and animal model experiments. However, the actual events in patients are quite different from the phenomena observed in in vitro studies. For example, short-term hypoxia in in vitro experiments is definitely different from chronic intermittent hypoxia in tumour tissues[9]. Intermediate and long-term intermittent hypoxia in tissues in vivo has a pressure screening effect, which can promote a distinctive molecular profile that is different from that in vitro [10]. In addition, hypoxia in vivo interacts with other microenvironments (e.g. extracellular matrix, inflammatory cells)[11, 12]. Therefore, it is necessary to investigate the role of hypoxia in tumours at the in vivo and tissue levels.

Because hypoxia exposure can cause changes in gene expression levels, researchers have begun to use hypoxia-related gene signatures to reflect the hypoxia of tumour tissues. For example, Malenstein et al. developed a 7-gene set associated with chronic hypoxia exposure that can be used to indicate the prognosis of HCC patients [13]. However, most previous studies only built hypoxia-related signatures and only focused on the relationship between the signatures and patient prognosis but did not reveal the panoramic view of molecular changes induced by hypoxia at the tissue level and especially in large cohorts. With the development and popularization of high-throughput technology, especially The Cancer Genome Atlas (TCGA) project, which has been gradually completed, increasingly more well organized genomic, epigenomic, transcriptomic and proteomic data have become available. This provides an unprecedented opportunity for an in-depth exploration of roles and mechanisms of hypoxia in in vivo tissues. Recent comprehensive studies of pan-cancer based on hypoxia-related gene signatures revealed some molecular landmarks of tumour hypoxia across cancer types at the tissue level, including some interesting and enlightening data on HCC [14-16]. However, the signatures used in the aforementioned studies are not HCC-specific, and because they are pan-cancer studies, the coverage and depth of the research content for HCC are limited. To solve this issue, we developed a novel, more robust HCC-specific hypoxia gene signature that can better reflect hypoxia exposure in HCC tissues. We performed a molecular classification of HCC using the signature and explored the relationship between the signature and patient prognosis. More importantly, we used this signature to comprehensively explore changes in hypoxia-related molecular features and changes in the immune microenvironments in HCC from genomic, epigenomic, transcriptomic, and proteomic perspectives. Our evidence suggests that hypoxia may play a more important role in the development of HCC than expected. We believe that the proposed hypoxia gene signature and the molecular landscapes revealed by it will provide useful information for the diagnosis and treatment of HCC.

## Method

### Cell culture and hypoxia treatment

HCC cell lines including HUH7, SNU-182 and HLF cells were used in present study. HUH7 and HLF cells from the Japanese Cancer Research Resources Bank. SNU-182 cells were obtained from American Type Culture Collection. HUH7 and HLF cells were cultured in Dulbeccos Modified Eagle Medium (DMEM, high glucose). SNU -182 were cultured in RPMI 1640 Medium. All medium was supplemented with 10% fetal calf serum (FCS) and 1% Penicillin (100 IU/m)-Streptomycin (100 g/ml) solution. Cells were maintained in an incubator with 37 □C, 5 %CO_2_, and 95% relative saturation of humidity. For hypoxia treatment, cells were cultured under an atmosphere of 1% O_2/_5% CO_2_/94% N_2_ for 24 h.

### RNA extraction and microarray

Hypoxia treated HUH7, SNU-182 and HLF cells were collected. Total RNA was extracted from treated cells using TRIzol Reagent following the protocol from manufacturers instructions. The concentration and quality of total RNA were measured using Nano Drop microvolume spectrometer. The RNA samples with high quality were labeled and hybridized on Agilent Whole human genome chip (4×44K). Differential expressed mRNAs were extracted using R software (version 3.6.1) with Limma package.

### Generation of a novel hypoxia gene signature and calculation of hypoxia score

Our novel hypoxia gene signature were built using robust rank aggregation(RRA) algorithm[17] from the microarray data of hypoxia treated HUH7, SNU-182 and HLF cells. Up-regulated (fold change >2) mRNAs with P<0.01 were selected from the RRA output. Totally 21 mRNAs were included. To estimate the hypoxia exposure, hypoxia score was calculated using gene set variation analysis (GSVA)[18]. We also compared our 21-gene hypoxia signature with seven other published hypoxia gene signatures including Buffa’s signature[19],Eustace’s signature[20],Ragnum’s signature[21], Sorensen’s signature[22], winter’s signature[23] and Malenstein’s signature[13]. Hypoxia score based these seven signatures were also calculated by GSVA.

### Public datasets in GEO

We retrieved three independent mRNA microarray datasets (GSE18494, GSE55214 and GSE57613) based on hypoxia treated HCC cells from the Gene Expression Omnibus (GEO, https://www.ncbi.nlm.nih.gov/geo/). Datasets and available clinical information of ten HCC cohorts (GSE14520, GSE22058, GSE25097, GSE36376, GSE45436, GSE64041, GSE76297, GSE76427, GSE10141 and GSE9843) were downloaded from GEO. All gene symbols in GEO datasets were converted to the latest (HUGO Gene Nomenclature Committee) HGNC Symbols.

### Multi-omic data and clinical data of TCGA-LIHC

mRNA expression, microRNA expression, lncRNA expression, CNAs, SNVs, DNA methylation data of the Cancer Genome Atlas Liver Hepatocellular Carcinoma (TCGA-LIHC) were downloaded from the TCGA data portal (https://portal.gdc.cancer.gov/). The RNA alternative splicing data of TCGA-LIHC were obtained from TCGA SpliceSeq (http://projects.insilico.us.com/TCGASpliceSeq/). The reverse phase protein array (RPPA) data of TCGA-LIHC were downloaded from the cancer proteome atlas (https://www.tcpaportal.org/tcpa/index.html).

### Biological process and pathway enrichment assay

The biological process and pathway enrichment assay of candidate genes were performed using online tools provided by Metascape (http://metascape.org/gp/index.html). The enrichment analysis has been carried out with the following ontology sources: KEGG Pathway, GO Biological Processes, Reactome Gene Sets, Canonical Pathways and CORUM. All genes in the genome have been used as the enrichment background. To establish the network based on the relationships between the terms, a subset of enriched terms with a similarity > 0.3 were connected by edges. 20 clusters were obtained and the terms with the best p-values were selected (more details see the website of Metascape). The network is visualized using Cytoscape (version 3.7.2).

### Protein-protein Interaction Enrichment Analysis

We used online tools provided by Metascape to perform protein-protein interaction enrichment analysis for the production of candidate genes. According to data from BioGrid, InWeb_IM and OmniPath, a resultant network contains the subset of proteins that form physical interactions with at least one other member were built using Molecular Complex Detection (MCODE) algorithm[24]. Then Pathway and process enrichment analysis has been applied to each MCODE component (more details see the website of Metascape). The network is visualized using Cytoscape (version 3.7.2).

### Gene set enrichment analysis (GSEA)

GSEA was performed for candidate mRNAs cross HCC cohorts using GSEA tools (version 4.0.3) provided by the Molecular Signatures Database (MSigDB, https://www.gsea-msigdb.org/gsea/msigdb/index.jsp). The hallmark gene sets as the reference gene set. The significantly activated or suppressed pathways were Identified as pathways with p value<0.05 and FDR<0.25.

### Interactions analysis for mRNA/microRNA/lncRNA

miRNA-target interactions were presented by intersecting the predicting target sites of miRNAs with binding sites of Ago protein using the Encyclopedia of RNA Interactomes (ENCORI, http://starbase.sysu.edu.cn/index.php) and miRwalk 3.0 (http://zmf.umm.uni-heidelberg.de/apps/zmf/mirwalk/). The miRNA-lncRNA interactions were presented using LncBase v.2 experimental module tools (http://diana.imis.athena-innovation.gr/DianaTools/index.php). Interactions of mRNA/microRNA/lncRNA were visualized using Cytoscape (version 3.7.2).

### Estimation for immune microenvironment

The presence of infiltrating stromal and immune cells in tumor tissues were predicted using ESTIMATE algorithm (Estimation of Stromal and Immune cells in Malignant Tumor tissues)[25]. ESTIMATE algorithm provided stromal score (that captures the presence of stroma) and immune score (that represents the infiltration of immune cells) based on mRNA expression of tumor tissues. The relative levels of distinct immune cell types were estimated using CIBERSORT tools (https://cibersort.stanford.edu) with LM22 files as reference.

### Statistical analysis

Statistical analyses were performed using R software (version 3.6.1) with relevant packages. In brief, the differential expressed mRNAs were extracted from microarray datasets using Limma package. The differential expressed mRNAs, microRNA and lncRNA of TCGA-LIHC were identified using Linnorm packages. The difference between two groups were compared using independent t test or Wilcox test. Adjusted P-Value were obtained using FDR (False Discovery Rate) method. Coefficients were calculated using Pearson or Spearman correlation analysis. Chi-squared test was used to determine the significant difference between the frequencies. Survival analysis were performed using Univariate Cox/multivariate analysis hazard analysis or Kaplan-Meier survival estimate using survival package. The forest-plot R package was employed to visualize the hazard rate obtained from survival analysis. The Kaplan-Meier survival curves were created using survminer package with Logrank test. A prognostic model based on hypoxia score were established using Lasso (least absolute shrinkage and selection operator)-cox Regression method. The model was visualized and validated using Hdnom package. In the present study, statistical significance was set at a probability value of P < 0.05

## Results

### Identification for a novel hypoxia signature that includes 21 genes

First, through microarray analysis, differentially expressed genes with fold changes (FC) satisfying log_2_FC > 0.58 or log_2_FC <-0.58 and *P* < 0.05 in HUH7, SNU-182 and HLF cells were obtained after 24 h of hypoxia exposure. Next, we started to construct a new hypoxia signature based on the data obtained according to a design principle that involved using a minimum number of genes to collectively indicate the hypoxia-related phenotype of interest. The differentially expressed genes of 3 hypoxia-treated HCC cell lines were screened and integrated using the robust rank aggregation (RRA) algorithm. After a literature review, 21 out of these differentially expressed genes with FC > 2 were selected to construct the hypoxia signature. The expression of these 21 genes was significantly increased in HUH7, SNU-182 and HLF cells after hypoxia exposure. We also found 3 hypoxia-related microarray datasets in the Gene Expression Omnibus (GEO) database, including mRNA expression in hypoxia-treated HEPG2 (GSE18494), HUH7 (GSE55214) and HEP3B (GSE57613) cells. In the 3 datasets, the 21 genes that were selected to construct the hypoxia signature were also significantly upregulated after hypoxia exposure (Figure 1a). Therefore, these 21 genes are hypoxia-responsive genes in HCC cells. Next, gene set variation analysis (GSVA) were used to calculate the hypoxia score. Compared with the control group, hypoxia-treated HCC cells had significantly higher hypoxia scores (Figure 1b). Therefore, the hypoxia scores calculated based on the 21-gene hypoxia signature can reflect the hypoxic condition in HCC cells. To further prove the robustness of the 21-gene hypoxia signature in the assessment of hypoxia, 6 hypoxia signatures that have been reported in highly cited articles were selected to calculate hypoxia scores: Buffas signature (15 genes), Eustaces signature (26 genes), Ragnums signature (32 genes), Sorensens signature (27 signature), Winters signature (99 genes) and Malensteins signature (7 genes). The first 5 have been proven to be excellent performers in recent pan-cancer comprehensive studies assessing the robustness of different hypoxia signatures[14, 15]. Malensteins signature is a liver -specific hypoxia signature associated with HCC prognosis[13]. There were significantly positive correlations between hypoxia scores calculated based on our 21-gene hypoxia signature and those calculated based on the other 6 signatures for cancer tissues from 9 HCC datasets, including The Cancer Genome Atlas-Liver Hepatocellular Carcinoma (TCGA-LIHC), GSE14520, GSE22058, GSE25097, GSE36376, GSE45436, GSE64041, GSE76297, and GSE76427 (Figure 1c). Cell-and tissue-level evidence suggests that our 21-gene signature is robust for the assessment of hypoxia. The hypoxia scores for liver cancer tissues were clearly clustered in 2 groups (Figure 1 d), positive and negative, suggesting that the degrees of exposure of liver cancer tissues to hypoxia in different patients are different.

**Figure 1.**
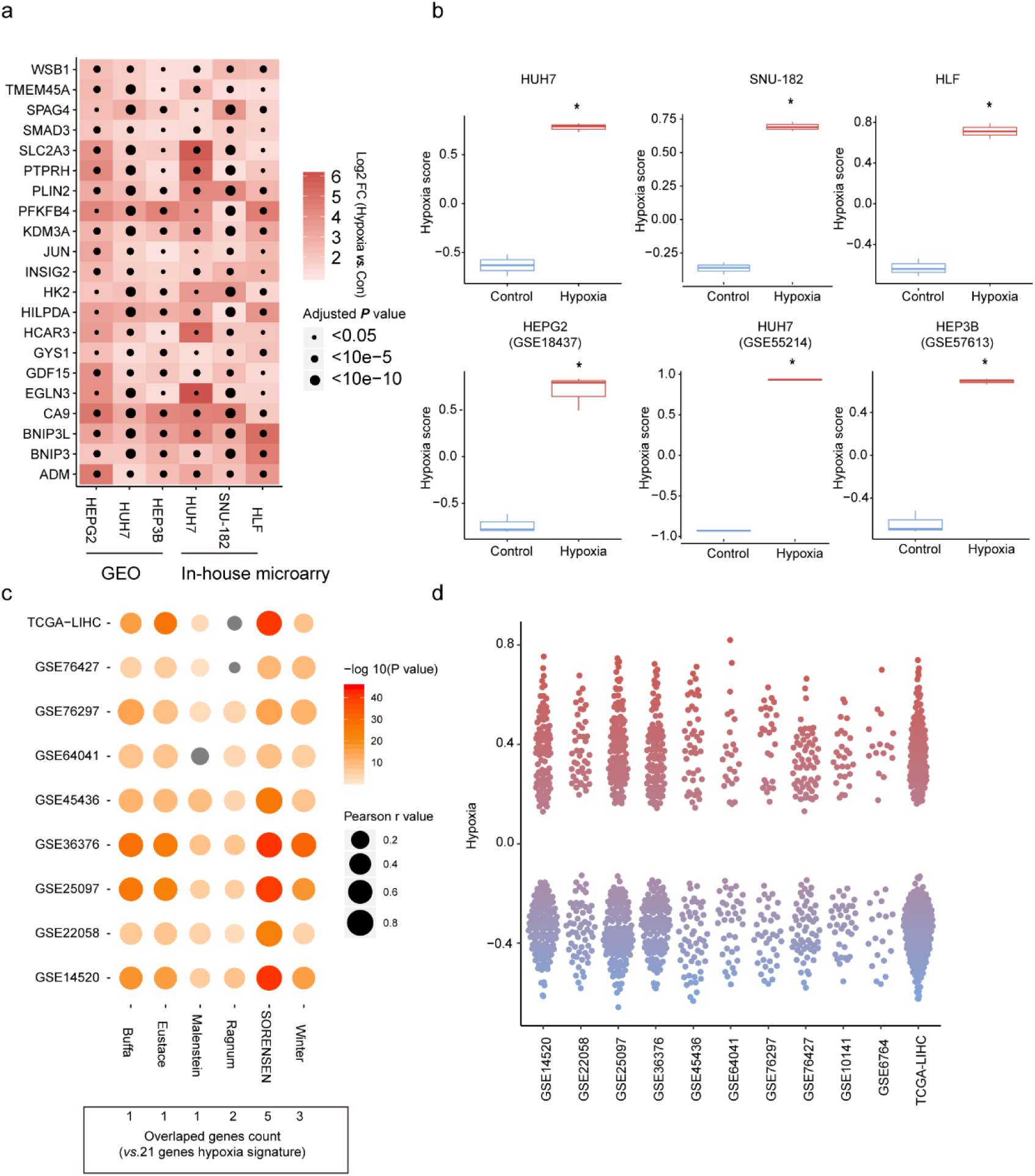
The 21-gene hypoxia signature indicated hypoxia exposure in hepatocellular carcinoma (HCC) cells. a) Expression changes in 21 genes in 6 HCC cells after hypoxia exposure. b) Hypoxia scores calculated based on the 21-gene hypoxia signature using gene set variation analysis (GSVA) significantly increased in hypoxia-treated HCC cells. c) Correlations between the hypoxia scores calculated based on the 21-gene hypoxia signature and the hypoxia score calculated based on other published hypoxia signatures in the cancer tissues of 9 HCC cohorts. d) The distribution of hypoxia scores calculated based on the 21-gene hypoxia signature in the cancer tissues of 11 HCC cohorts. * compared with the control group, *P* < 0.05.

### HCC subtypes obtained based on the 21-gene hypoxia signature have different clinical characteristics and prognoses

Unsupervised hierarchical clustering analysis was used to group patients (n = 374) in the TCGA-LIHC dataset based on the 21-gene signature. All patients were grouped into 2 subtypes based on the 21-gene signature (Figure 2a): LIHC-cluster A (n = 315) and LIHC-cluster B (n = 59). The proportion of patients with stage III-IV HCC, according to the tumour-node-metastasis (TNM) staging system, in LIHC-cluster B was higher (χ^2^ = 11.35, *P* < 0.01) than that in LIHC-cluster A, and the alfa-fetoprotein (AFP) level in LIHC-cluster B was higher than that in LIHC-cluster A (χ^2^ = 4.44, *P* < 0.05). The overall survival (OS) rate (Figure 2b, hazard ratio (HR) = 2.15, log-rank *P* < 0.01) and disease-free survival (DFS) (Figure 2c, HR = 2.17, log-rank *P* < 0.01) of patients in LIHC-cluster B were significantly lower than those of patients in LIHC-cluster A. The hypoxia scores of patients in LIHC-cluster B were significantly higher than those of patients in LIHC-cluster A (Figure 2d). Similar unsupervised hierarchical clustering analysis grouped the patients in GSE14520 into 3 subtypes (Figure 2e): GSE14520-cluster A (n = 41), GSE14520-cluster B (n = 35) and GSE14520-cluster C (n = 172). Compared with the other 2 subtypes, GSE14520-cluster A had a significantly higher proportion of patients with high TNM stages (χ^2^ = 15.07, *P* < 0.01) and higher AFP levels (χ^2^ = 7.54, *P* < 0.05). In GSE14520-cluster A, the number of cases with a tumour diameter > 5 cm was also higher than that in the other 2 subtypes (χ^2^ = 9.91, *P* = 0.01). Roessler et al. used a metastasis gene signature to group GSE14520 patients into patients with high invasion risk and patients with low invasion risk[26]. In GSE14520-cluster A, the proportion of patients with high invasion risk was significantly higher than in the other 2 subgroups (χ^2^ = 22.36, *P* < 0.01). As expected, the OS rate (Figure 2f, HR_A: B_ = 1.31, HR_A: C_ = 2.71, log-rank *P* < 0.01) and DFS (Figure 2G, HR _A: B_ = 1.81, HR _A: C_ = 1.94, log-rank *P* < 0.01) of patients in GSE14520-cluster A were lower than those of patients in other subtypes. Patients with a poor prognosis in GSE14520 cluster A had high hypoxia scores (Figure 2g). The evidence suggests that the molecular classification of HCC based on the 21-gene hypoxia signature can reflect different clinical features and prognoses.

**Figure 2.**
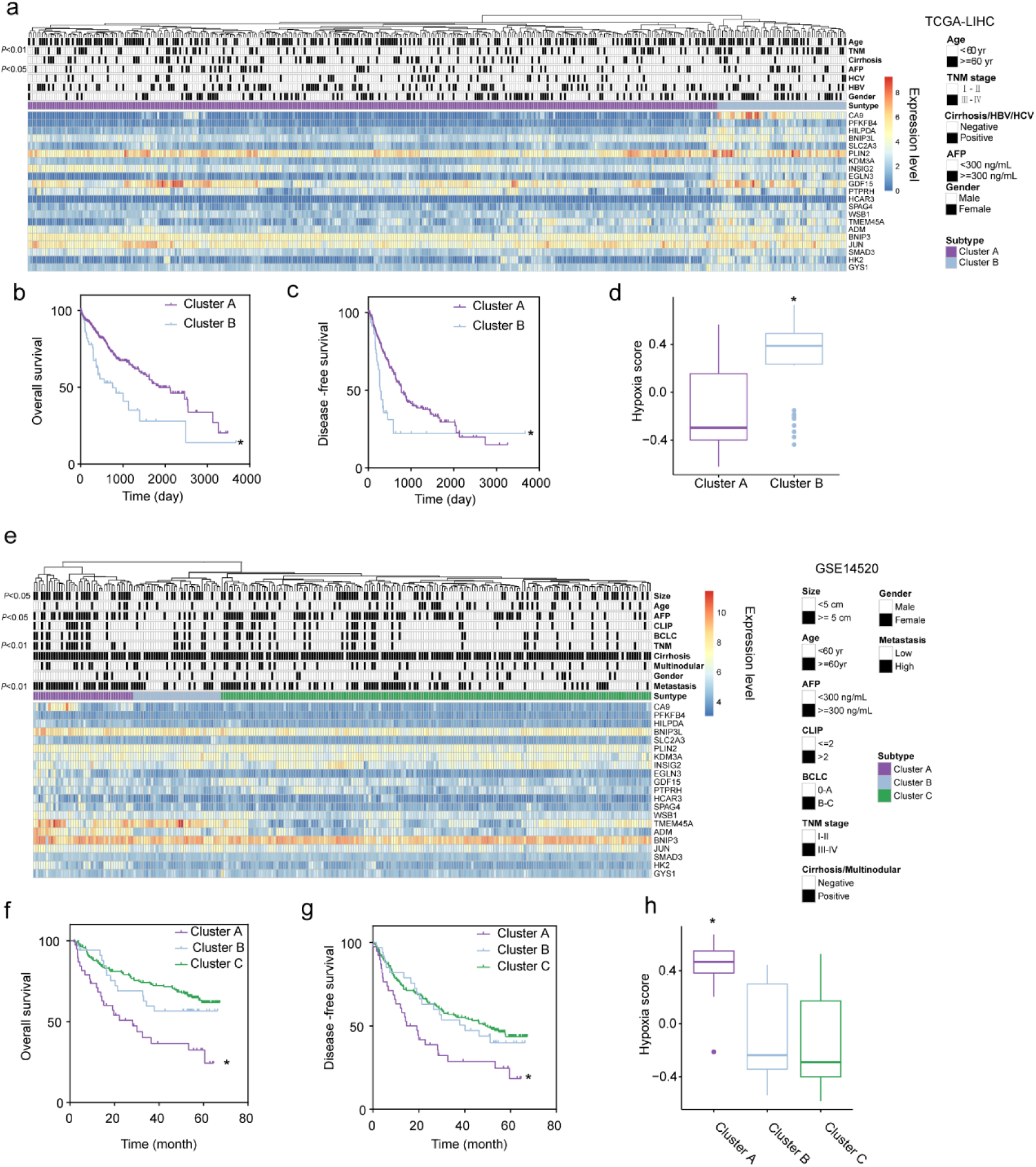
Hepatocellular carcinoma (HCC) subtypes classified using the 21-gene hypoxia signature are correlated with clinical characteristics and patient prognoses. a) Unsupervised clustering based on the 21-gene hypoxia signature grouped patients in TCGA-LIHC into 2 subtypes. There were significant differences in tumour-node-metastasis (TNM) stage and alfa-fetoprotein (AFP) levels between the 2 subtypes. b) and c) Differences in OS and DFS between the 2 subtypes in TCGA-LIHC. d) Difference in hypoxia scores between the 2 subtypes in TCGA-LIHC. e) Unsupervised clustering based on the 21-gene hypoxia signature grouped patients in GSE14520 into 3 subtypes. There were differences in tumour size, AFP levels, TNM stage, and invasiveness between cluster A and the other clusters. f) and g) Differences in OS and DFS between the 3 subtypes in GSE14520. h) Differences in hypoxia scores between the 3 subtypes in GSE14520. * compared with other clusters, *P* < 0.05.

### Hypoxia scores calculated based on the 21-gene hypoxia signature can be used as a prognostic marker that indicates HCC progression

The results of the univariate and multivariate Cox regression analyses indicate that a high hypoxia score is an independent risk factor for OS in HCC patients in the TCGA-LIHC and GSE14520 datasets (HR = 1.82 in TCGA-LIHC and HR = 2.71 in GESE14520, all *P* < 0.05). In TCGA-LIHC and GSE14520, the hypoxia score was used to evaluate the receiver operating characteristic (ROC) curves for 1-year, 3-year and 5-year survival and the corresponding areas under the curves (AUCs) (Figure 3a and b). The optimal cut-off value was calculated using X-tile software[27], and the patients were divided into a high hypoxia score group and a low hypoxia score group. The OS rate for patients in the high hypoxia score group was lower than that for patients in the hypoxia score group. After stratification of patients using TNM staging, a high hypoxia score still implied poor OS (Figure 3c and d). In the TCGA-LIHC and GSE14520 datasets, patients in the high hypoxia score group had shorter DFS than did patients in the low hypoxia score group (Figure 3e). The TCGA-LIHC dataset was used as the training set, and TNM stages and hypoxia scores were used as factors to construct the least absolute shrinkage and selection operator (LASSO)-Cox regression model (c-index = 0.73) for predicting the OS of patients, and a predictive nomogram was established accordingly (Figure 3f). The effect of the model was verified in the GSE14520 dataset. In another HCC dataset, GSE76427, patients with a high hypoxia score also had a low OS rate (Figure 3g). Therefore, the hypoxia score calculated based on the 21-gene signature has the potential to be a prognostic marker for HCC patients. Compared with the low hypoxia score group in the TCGA-LIHC dataset, the high hypoxia score group (≥ median hypoxia score) had a higher number of cases with late TNM stages. In the GSE14520 dataset, patients in the high hypoxia score and low hypoxia score groups differed significantly in AFP level, degree of cirrhosis, tumour size, TNM stage, Barcelona Clinic Liver Cancer (BCLC) stage, Cancer of the Liver Italian Program (CLIP) stage, and multinodular status, suggesting that a high hypoxia score is an unfavourable factor for HCC patients and can indicate progression. In the GSE6764 dataset, the hypoxia score in HCC tissues gradually increased from very early to very advanced HCC (Figure 3h). Metastasis is an important step in the progression of HCC. Roessler and Chen independently used 2 different signatures to group patients in the GSE14520 dataset as having a high metastasis risk or a low metastasis risk[26, 28]. We found that patients with a high metastasis risk had higher hypoxia scores, suggesting that hypoxia scores can indicate the risk of metastasis in HCC patients (Figure 3i). The data from the TCGA-LIHC and GSE10141 datasets suggest that high hypoxia scores can indicate vascular invasion (Figure 3j).

**Figure 3.**
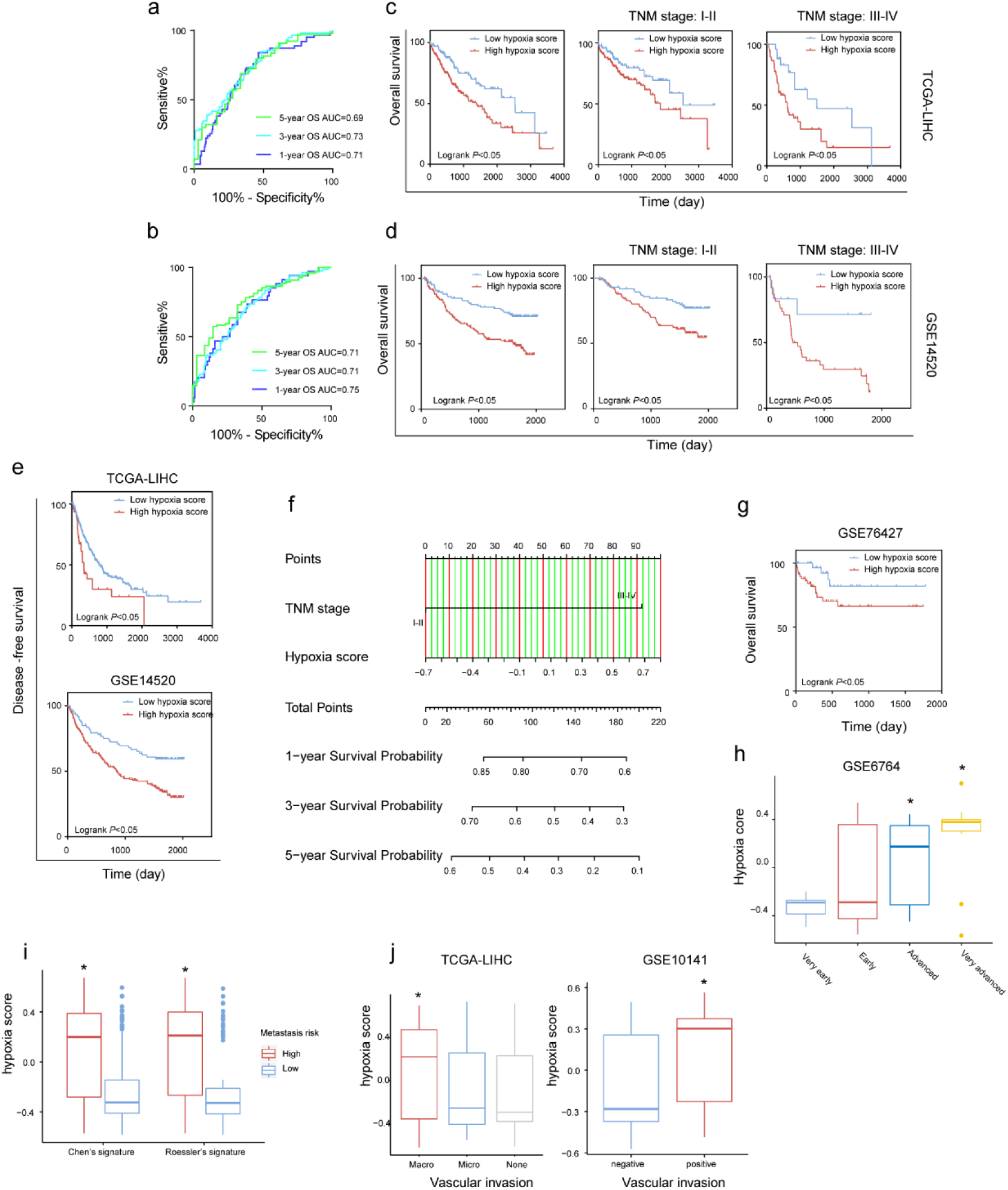
The hypoxia scores calculated based on the 21-gene hypoxia signature can predict the prognoses of patients with hepatocellular carcinoma (HCC). a) and b) The time-dependent receiver operating characteristic (ROC) curves for overall survival (OS) of patients in TCGA-LIHC and GSE14520 estimated by hypoxia scores. c) and d) The differences in OS between patients with high hypoxia scores and low hypoxia scores in TCGA-LIHC and GSE14520. After stratification of patients using TNM staging, the differences in OS between patients with high hypoxia scores and low hypoxia scores were compared again. The cut-off was calculated using X-tile software. e) The differences in DFS between the patients with high hypoxia scores and low hypoxia scores in TCGA-LIHC and GSE14520. f) Nomogram of the LASSO-cox regression model to predict 1-, 3-or 5-year OS in TCGA-LIHC using hypoxia score and TNM stage. g) The difference in OS between patients with high hypoxia scores and low hypoxia scores in GSE76427. The cut-off value was calculated using X-tile software. h) The hypoxia scores for patients in GSE6764 increased with the progression of clinical stage. i) The differences in hypoxia scores between HCC patients with different metastasis risk. j) Correlations between hypoxia scores and vascular invasion in patients in TCGA-LIHC and GSE10141. * compared with other groups, *P* < 0.05.

### Transcriptomic alterations in HCC patients with different hypoxia scores

First, we analysed the differences in mRNA expression between HCC patients with high hypoxia scores (greater than the upper quartile) and those with low hypoxia scores (less than the lower quartile). Here, mRNA with log_2_FC > 0.58 or log_2_FC <-0.58 and adjusted *P* < 0.05 was defined as differentially expressed mRNA (DE-mRNA). The genes in the 21-gene signature are marked in the volcano plot. The members of the 21-gene signature in the TCGA-LIHC and GSE14520 datasets were almost all upregulated in HCC patients with high hypoxia scores. After determining similar DE-mRNAs in the TCGA-LIHC and GSE14520 datasets, the integrated DE-mRNA list (341 mRNA) was subjected to functional enrichment analysis and Cox survival analysis.

We also integrated DE-mRNA data for 11 HCC cohorts. In these cohorts, the proportion of DE-mRNA in the total mRNA measured was positively correlated with the interquartile range (IQR) of the liver cancer tissue hypoxia scores in this cohort (Figure 4a), suggesting that the difference in mRNAs was to some extent caused by the difference in hypoxia scores. Because the number of probes in the GSE10141 dataset was small, this dataset was excluded in subsequent studies. We counted the frequency of each mRNA identified as a DE-mRNA in all cohorts and included DE-mRNAs with a frequency equal to or greater than 5 in the high frequency/DE-mRNA (HF/DE-mRNA) list. The mRNA changes in this list were relatively consistent among the 10 cohorts. A total of 371 mRNAs were selected, including 192 upregulated DE-mRNAs and 179 downregulated DE-mRNAs. Based on the survival data from the TCGA-LIHC and GSE14520 datasets, we performed survival analysis on 371 HF/DE-mRNAs (log-rank test, cut-off = median expression level). Through Venny analysis, we integrated the survival data from the TCGA-LIHC and GSE14520 datasets and obtained 129 HF/DE-mRNAs related to HCC survival (log-rank P_TCGA-LIHC_ & log-rank P_GSE14520_ < 0.05), which included 59 risk factors (HR_TCGA-LIHC_ & HR_GE14520_ > 1) and 70 protective factors (HR_TCGA-LIHC_ & HR_GE14520_ <1). There were 51 HF/DE-mRNAs that met the log-rank P < 0.01 and HR < 0.7 or > 1.3 requirements in both of the datasets (TCGA-LIHC and GSE14520). All risk factors for OS were significantly upregulated in the high hypoxia score group while the protective factors were downregulated; this phenomenon occurred in almost all cohorts (Figure 4 b). For example, ANXA5 and SPP1 were both risk factors for OS in the TCGA-LIHC and GSE14520 datasets and were significantly overexpressed in the high hypoxia score group in 9 cohorts.

**Figure 4.**
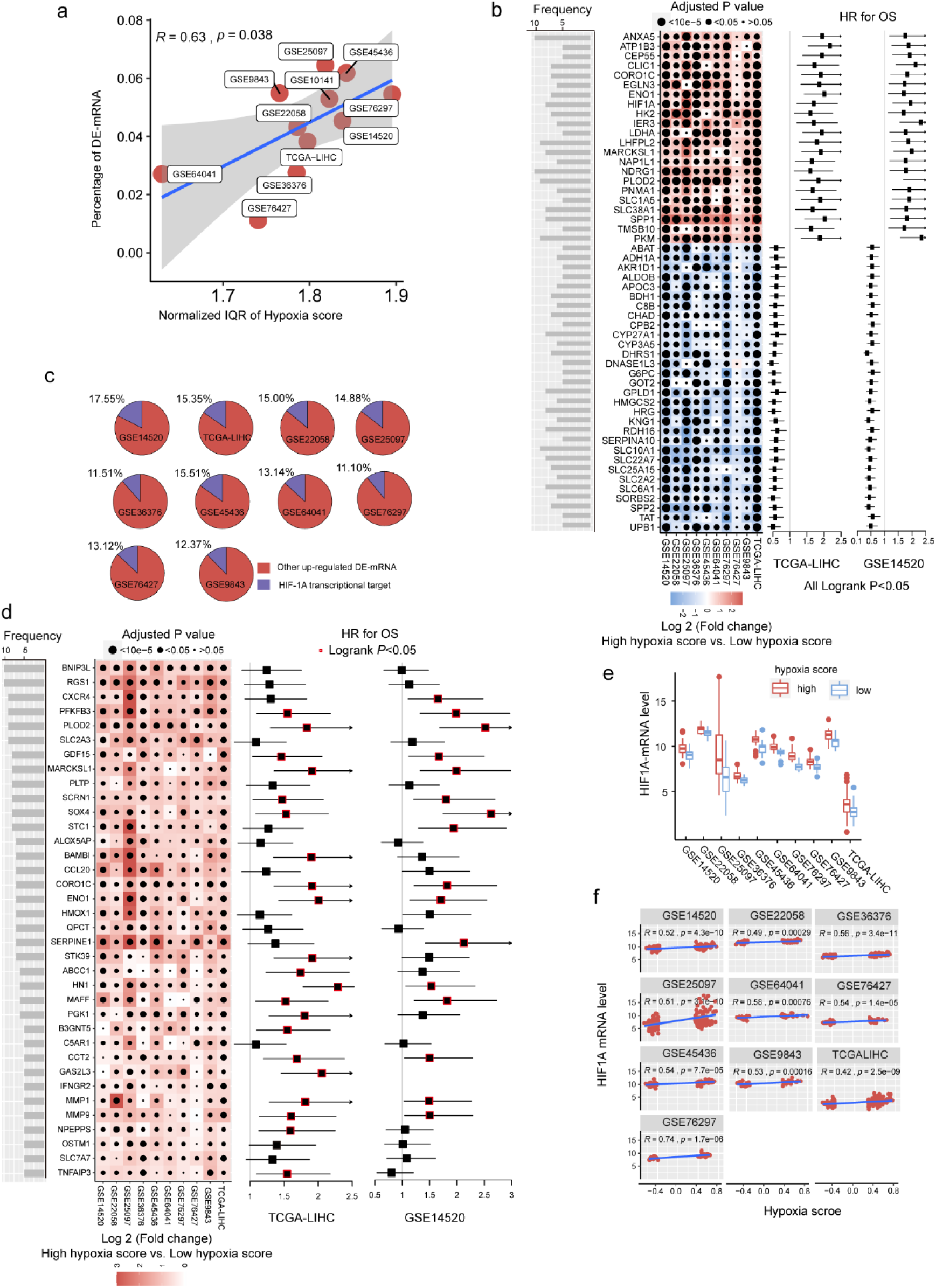
The mRNA alterations in hepatocellular carcinoma (HCC) patients with high hypoxia scores and low hypoxia scores. a) Hypoxia scores calculated based on the 21-gene hypoxia signature. According to the upper quartile and the lower quartile, patients were divided into a high hypoxia score group and a low hypoxia score group. In the 11 HCC cohorts, the percentage of differentially expressed (DE)-mRNAs among all mRNAs measured was positively proportional to the interquartile range (IQR) of the hypoxia scores. b) A total of 129 high frequency/DE-mRNAs (HF/DE-mRNAs) are correlated to HCC patient survival. The heat map shows the difference in the expression of these mRNAs between the high hypoxia score group and the low hypoxia score group in the 10 HCC cohorts, that is, the log_2_ (fold change) between the 2 groups. The forest plot indicates the hazard ratios (HRs) of these mRNAs for OS in the survival analysis (all log-rank *P* < 0.01, HR < 0.7 or > 1.3, cut-off value = median expression level). c) The percentage of transcription targets with differentially expressed hypoxia-inducible factor 1-alpha (HIF-1A) in a dataset for all DE-mRNAs in the dataset. d) Thirty-six mRNAs may function as transcription targets of HIF-1A, and the upregulation trends are consistent in the 10 HCC datasets. The heat map shows the difference in the expression of these mRNAs between the high hypoxia score group and the low hypoxia score group. The forest plot indicates the HRs of these mRNAs for OS in the survival analysis (cut-off = median expression level). e) The differences in HIF-1A mRNA expression levels between the high hypoxia score group and the low hypoxia score group in 10 HCC datasets. f) Correlations between HIF-1A mRNA expression levels and hypoxia scores for the 10 HCC datasets.

Hypoxia-inducible factor 1-alpha (HIF-1A) plays a core role in hypoxia, and 10%-15% of the upregulated DE-mRNAs in each cohort may be potential transcription targets of HIF-1A (Figure 4c). Among these potential HIF-1A transcription targets, DE-mRNAs with a frequency higher than 5 in each cohort are shown in Figure 4d; most are risk factors for survival. In addition, although HIF-1A protein levels are known to be regulated after translation under hypoxic conditions, 10 HCC datasets indicated that the mRNA level of HIF-1A significantly increased in the high hypoxia score group and showed a significantly positive correlation with hypoxia score (Figure 4e and f). This suggests that in addition to HIF-1A protein level regulation under hypoxic conditions, the change in HIF-1A mRNA level is also worthy of attention.

To further reveal the functions of HF/DE-mRNAs, the enrichment of biological processes and pathways involving HF/DE-mRNA were analysed using data from different sources, such as Gene Ontology (GO) biological processes, Kyoto Encyclopedia of Genes and Genomes (KEGG) pathways, Reactome Gene Sets, and canonical pathways (Figure 5a). In addition to the response to hypoxia, HF/DE-mRNAs are mainly involved in biological processes related to metabolism, including glucose metabolism, lipid metabolism and amino acid metabolism. As expected, pathway enrichment analysis results for data from multiple sources all showed significant enrichment of the HIF-1 pathway and glucose metabolism pathways; other enriched pathways were mainly related to various metabolic pathways. The pathways related to the regulation of the extracellular matrix were also associated with HF/DE-mRNAs. Therefore, we hypothesized that hypoxia exposure might affect the extracellular matrix, which determines tumour invasion and metastasis. To find the connection between terms in the enrichment analysis of biological processes and pathways, we clustered the terms and constructed a network (Figure 5b). The name of each cluster is the name of the most representative term, and the node size is the number of genes in the term.

**Figure 5.**
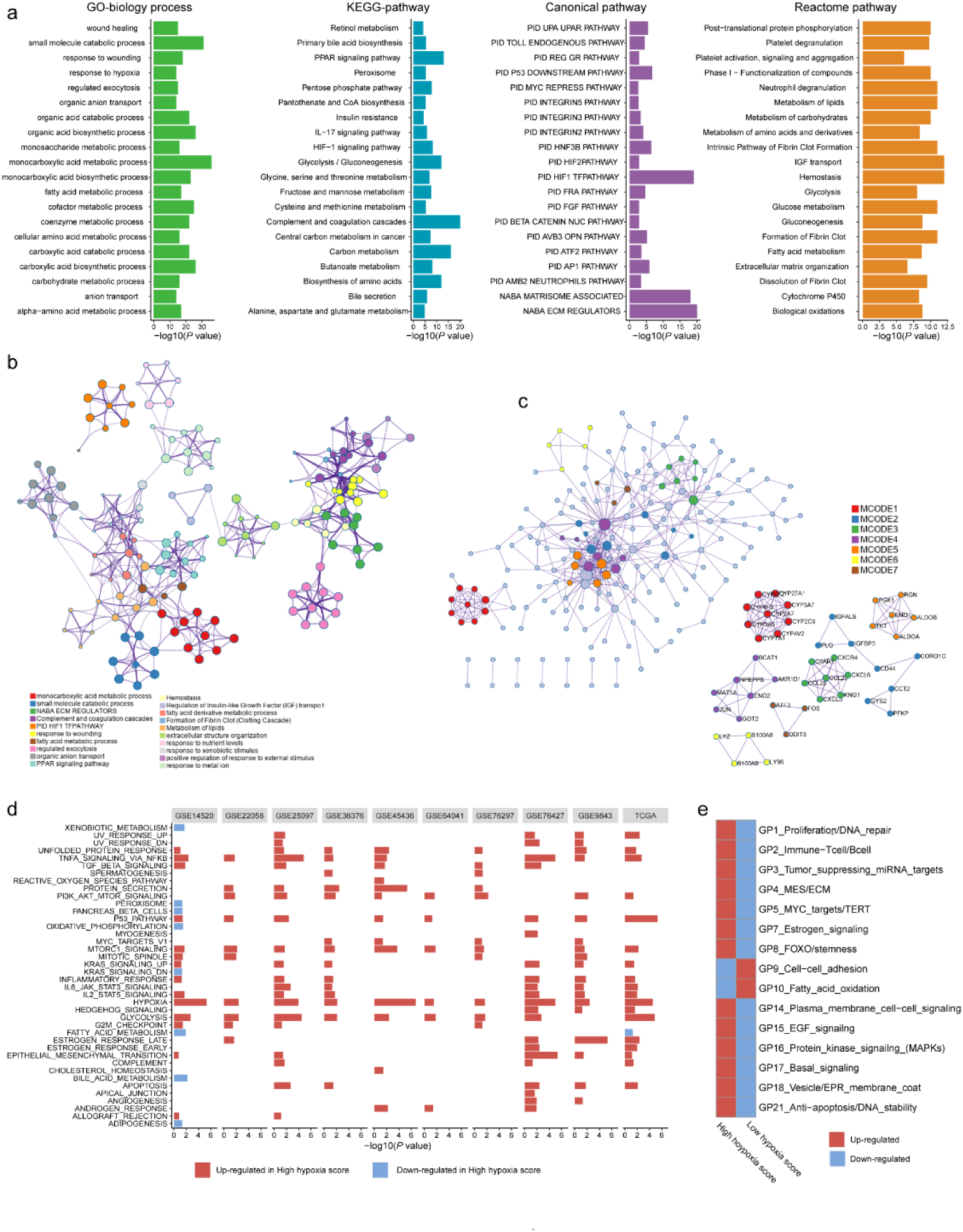
Biological processes and pathway functional enrichment analysis of 371 high frequency/DE-mRNAs (HF/DE-mRNAs) extracted from 10 hepatocellular carcinoma (HCC) datasets. a) The top 20 (sorted by *P* value) from each enrichment analysis result for GO biological processes of HF/DE-mRNAs and the pathway enrichment analysis results for 3 data sources. b) A clustering network formed by correlated terms from the functional enrichment analysis. c) HF/DE-mRNA translation products constructed based on the molecular complex detection (MCODE) algorithm. d) Protein-protein interaction (PPI) enrichment network. d) Gene set enrichment analysis (GSEA) of HF/DE-mRNAs in 10 datasets, showing the pathways with *P* < 0.05 and false detection rate (FDR) <0.25. The reference gene sets are the hallmark gene sets. In the TCGA-LIHC dataset, the differences between the high hypoxia score group and the low hypoxia score group in 15 gene programmes related to biological behaviours of tumours. These gene programmes were identified by Hoadleys team, and the upregulation and downregulation trends were calculated as single-sample GSEA z-scores.

For the HF/DE-mRNA translation products, protein-protein interaction (PPI) enrichment analysis was carried out with the following databases: BioGrid6, InWeb_IM7, and OmniPath8. We constructed a PPI enrichment network of physical interactions using molecular complex detection (MCODE) (Figure 5c). In addition to focusing on HF/DE-mRNAs, we used gene set enrichment analysis (GSEA) to reveal differences between hallmark gene sets between the high hypoxia score group and low hypoxia score group based on all mRNA differences between the 2 groups (Figure 5d). The hypoxia gene set was upregulated in all cohorts with high hypoxia scores. The genes composing the glycolysis gene set was upregulated in the high hypoxia score groups of 9 cohorts. Other upregulated gene sets with high consistency in the 10 cohorts included the P53 pathway, PI3K/AKT/mammalian target of rapamycin (mTOR) signalling, TNFA signalling via NFKB, unfolded protein response, TGF-beta signalling, and MTORC1 signalling. The activation of these pathways is critical to the onset of progression of tumours and can be used to explain the poor prognosis of patients in the high hypoxia score group. In addition, many drugs target these pathways, suggesting that the degree of hypoxia exposure needs to be considered in the selection of medications for patients with HCC. Although the fatty acid metabolism pathway was downregulated in the high hypoxia score groups in the TCGA-LIHC and GSE14520 cohorts, downregulation was not consistent among the other pathways. After applying a bimodality filter and weighted gene correlation network-based clustering, Hoadleys team identified 22 nonredundant gene programmes related to biological behaviours of tumours[29]. We found 15 gene programmes that were significantly different between the high hypoxia score group and the low hypoxia score group (Figure 5e), and the single-sample GSEA z-scores for some cancer-promoting gene programmes increased in patients with high hypoxia scores.

We used TCGA-LIHC data to analyse the differences in microRNAs (miRs) in HCC tissues with high hypoxia scores and low hypoxia scores. The miRs with log_2_FC > 0.58 or log_2_FC < -0.58 and adjusted *P* < 0.05 were defined as differentially expressed miRs (DE-miRs). We found a total of 63 DE-miRs, including 39 upregulated DE-miRs and 24 downregulated DE-miRs. Survival analysis showed that some DE-miRs were related to the OS rate (log-rank test, cut-off = median expression level) of HCC patients (Figure 6a). Among these DE-miRs, miR-210 is a frequently reported master hypoxia-responsive miR, and its high expression has been observed in a variety of hypoxia-treated tumour cells[30, 31]. In our study, miR-210-3p had the smallest adjusted *P* value among the upregulated DE-miRs (log_2_FC = 2.41). Survival analysis using the median expression level as the cut-off showed that miR-210-3p had the largest HR (log-rank *P* < 0.05) and that high miR-210-3p expression indicated a poor prognosis. Additionally, the passenger strand of miR-210 (miR-210-5p) was significantly upregulated in HCC tissues with high hypoxia scores, and high miR-210-5p expression also indicated a poor prognosis. The presence of miR-210 in the DE-miR list means that our 21-gene hypoxia signature can reflect hypoxia exposure in HCC. Among the downregulated DE-miRs, miR-139-5p had the smallest adjusted *P* value (log_2_FC = -0.68). The response of miR-139-5p to hypoxia has not been reported. We found that low miR-139-5p expression indicates poor outcomes of HCC patients. Next, we predicted target mRNAs of DE-miRs. Combined with the HF/DE-mRNAs list, we obtained 2 independent DE-miRs-HF/DE-mRNAs networks, including upregulated DE-miRs/downregulated DE-mRNAs and downregulated DE-miRs/upregulated DE-mRNAs. These 2 networks suggested that the changes in some DE-mRNAs in HCC tissue under hypoxic conditions might be caused by the targeted regulation of DE-miRs. For example, miR-216b-5p is the miR with the highest degree of reduced expression in the high hypoxia score group (log_2_FC = -2.61, adjusted *P* < 0.001), and it is complementary to the 3 untranslated region (3 -UTR) of HK2. The prediction was supported by Argonaute-crosslinking immunoprecipitation sequencing (AGO-CLIP-seq) data [32, 33]. HK2 is a definite hypoxia-inducible gene[34]. The above results suggest that the high expression of HK2 in hypoxia may be associated with the reduction in miR-216b-5p. These 2 networks provide additional explanations for hypoxia-induced transcriptome changes in HCC tissues in addition to the HIF-1A-related mechanism. Attention was paid to miRs with high connectivity in the network, such as miR-1224-5p, miR-877-5p, let-7a-2-3p, and miR-378c. HIF-1A mRNA has a targeted relationship with some downregulated miRs, such as miR-101-3p and miR-194-3p. These miRs can provide an explanation for the increase in HIF-1A mRNA expression in the high hypoxia score groups of the 10 cohorts. A survival-related refined network was obtained by combining the survival analysis results (Figure 6b). In this refined DE-miR/DE-mRNA network, all nodes were associated with the survival rate of HCC patients (log-rank *P* < 0.05), and a negative correlation between nodes was indicated (r < -0.3 and *P* < 0.05). For example, miR-194-5p showed low expression in the high hypoxia score group, which was a protective factor for the survival rate. Its potential target genes, SOX4, HK2, MARCKS, and LHFPL2, showed high expression in the high hypoxia score group and were risk factors for the OS rate. The expression of miR-194-5p was significantly negatively correlated with the mRNA expression of SOX4, HK2, MARCKS, and LHFPL2. The elements in the refined DE-miR/DE-mRNA network should receive more attention. We performed KEGG pathway enrichment analysis on all target genes of DE-miRs. The enrichment results of the top 20 (according to the P value) are shown in Figure 6c. Some important classical pathways related to tumour development, such as the Hippo pathway, TGF-beta pathway, Ras signalling pathway, mTOR pathway, PI3K-AKT pathway, Wnt pathway and AMPK pathway, were involved. This result further suggests that these DE-miRs may have an unignorable role in HCC tissues with high hypoxia scores. In addition, more than 80% (54/63) of the DE-miRs had at least 1 target gene enriched in the HIF-1 signalling pathway. This confirms that DE-miRs identified by the 21-gene hypoxia signature are indeed hypoxic-related.

**Figure 6.**
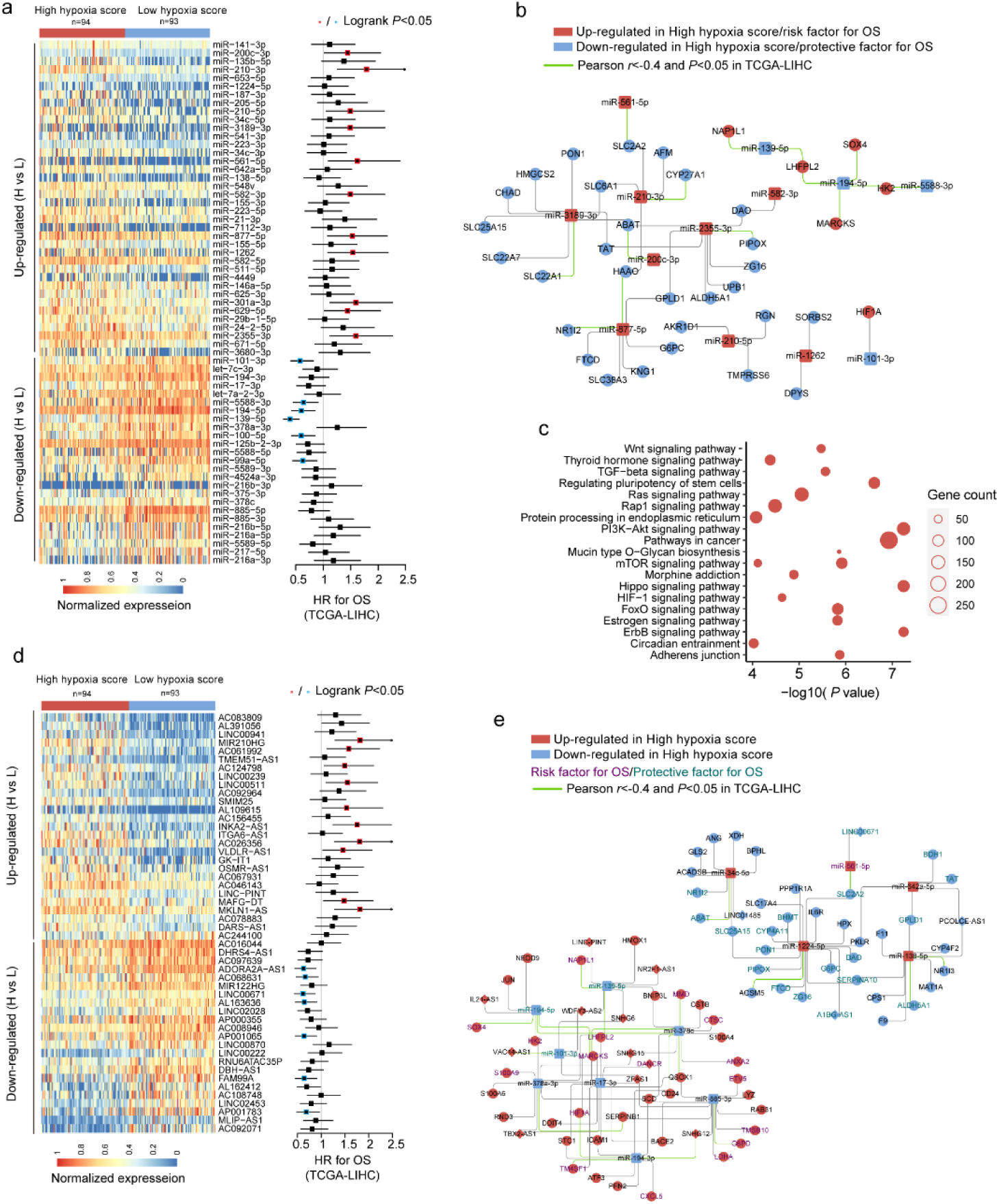
miRNA and long non-coding RNA (lncRNA) alterations in HCC patients with high hypoxia scores and low hypoxia scores. a) A total of 63 DE-miRNAs were significantly upregulated or downregulated in the high hypoxia score group. The forest plot indicates the hazard ratios (HRs) of these miRNAs for overall survival (OS) in the survival analysis (log-rank test). b) Some DE-miRNAs and HF/DE-mRNAs constitute a survival-related target interaction network. All nodes in the network are correlated with HCC patient survival in TCGA-LIHC (log-rank *P* < 0.05, cut-off = median expression level). The correlations between the nodes were calculated using Pearson correlation analysis. c) Top 20 (sorted by *P* value) KEGG pathway enrichment analysis results from 633 DE-miRNA target genes. d) Top 50 (sorted by adjusted *P* value) DE-lncRNAs that were significantly upregulated or downregulated in the high hypoxia score group. The forest plot indicates the hazard ratios (HRs) of these lncRNAs for OS in the survival analysis (log-rank test). e) The refined DE-lncRNA–DE-miRNA–HF/DE-mRNA ceRNA network. The correlations between nodes were calculated by Pearson correlation analysis. The survival data was from TCGA-LIHC. The cut-off is the median expression level.

Through the 21-gene signature, we revealed the presence of long non-coding RNAs (lncRNAs) that respond to and influence hypoxia exposure. In HCC tissues of patients with high hypoxia scores and low hypoxia scores, we found 719 differentially expressed lncRNAs (DE-lncRNAs), including 499 upregulated DE-lncRNAs and 220 downregulated DE-lncRNAs. The top-50 DE-lncRNAs (sorted by adjusted *P*) and survival analysis results (log-rank test, cut-off = median expression level) are shown in Figure 6d. miR210HG exhibited the most significant change (log_2_FC = 2.21) among the upregulated DE-lncRNAs. Higher miR210HG suggested poorer survival *(HR = 1.82*, log-rank *P <* 0.05). Similarly, lncRNAs AC124798 and AC061992 were also upregulated in the high hypoxia score group and suggested poor prognosis. In addition, lncRNAs LNC00671 and FAM99A were downregulated in the high hypoxia score group, which might be protective factors for prognosis. Based on competing endogenous RNA (ceRNA) theory and expression changes, we constructed a DE-lncRNA-DE-microRNA-HF/DE-mRNA network (Figure 6e). The relationship between each node and the survival rate for HCC patients was identified, and a negative correlation between nodes was also indicated. The miR-lncRNA relationship in the network is supported by experimental evidence (provided by DIANA-LncBase). Taking lncRNA-SNHG12 as an example, it exhibits high expression in the high hypoxia score group and is a risk factor for the survival of HCC patients. lncRNA-SNHG12 and miR-194-3p may have sequence complementarity. miR-194-3p was low in the high hypoxia score group and thus is a protective factor for HCC patient survival. The target mRNAs of miR-194-3p were TM4SF1 and HIF-1A. TM4SF1 and HIF-1A showed high expression in the high hypoxia score group and thus were risk factors for HCC patient survival. TM4SF1, HIF-1A and SNHG12 showed a significantly negative correlation with miR-194-3p. Therefore, a hypoxia-responsive lncRNA-SNHG12/miR-194-3p TM4SF1 or HIF-1A ceRNA network is likely present in the cancer tissues of HCC patients, and the ceRNA network is involved in tumour development and is related to patient prognosis. Leaving out sequence complementarity, we constructed the co-expression networks of all HF/DE-mRNAs, DE-miRs and DE-lncRNAs (|Pearson r| > 0.8 and *P* < 0.05), including a positive co-network and a negative co-expression network; the hub genes in the 2 networks were SERPINC1 and PKM, respectively. SERPINC1 was significantly downregulated in the high hypoxia score groups of 6 cohorts while PKM was significantly upregulated in the high hypoxia score groups of 10 cohorts. Compared with those in normal tissues, SERPINC1 was significantly lower and PKM was significantly higher in HCC tissues. Combined with the survival analysis results, we speculate that SERPINC1 and PKM play important roles, namely, cancer-suppressing and cancer-promoting functions, respectively, under hypoxia exposure. Among miRs, only microRNA-194-3p and microRNA-194-5p had a significantly negative correlation with HIF-1A. These data may help to explain the differences in HIF-1A mRNA between groups with high hypoxia scores and reflect the core role of HIF-1A mRNA. In summary, there were differences in the expression of mRNAs, miRs and lncRNAs between the high hypoxia score and low hypoxia score groups. Many cancer-promoting RNA molecules were upregulated in the high hypoxia score group while cancer-suppressing RNA molecules were downregulated in the high hypoxia score group. Some DE-mRNAs and/or DE-lncRNAs and DE-mRNAs may form regulatory networks that participate in the development of HCC and affect the prognosis of HCC patients.

### Genomic alterations in HCC patients with different hypoxia scores

TCGA-LIHC data were used to reveal somatic copy number aberrations (CNAs) and somatic single-nucleotide variants (SNVs) in HCC patients with different hypoxia scores. First, from the overall differences in gene-level CNAs in patients with high hypoxia scores (greater than the upper quartile) and low hypoxia scores (less than the lower quartile), we found that CNAs of approximately 13.7% (3396/24769) of the genes were concentrated in the high hypoxia score group. CNA events in 71 cancer genes (according to the definition of the Precision Oncology Knowledge Base (OncoKB) cancer gene list) were significantly differentially distributed between the 2 groups (Figure 7a). The copy number gain to copy number loss ratios for most genes were significantly increased in the high hypoxia score group. The roles and CNA tendencies (according to the proportions of homozygous deletions, single copy deletions, low-level copy number amplification and high-level copy number amplification) of these 71 cancer genes are provided in Figure 8a. For example, CDK4 is an oncogene, and its CNAs in the high hypoxia score group are mostly copy number gains. IRF1 is a tumour suppressor gene, and its CNAs in the high hypoxia score group are mostly copy number losses. The occurrence frequency of CNAs of some cancer genes was high in patients in the TCGA-LIHC cohort. These data suggest that the genomic instability of HCC patients may be related to hypoxia exposure. Next, we analysed the difference in gene-level SNVs between the 2 groups of patients. Unfortunately, we did not obtain much evidence that indicated a strong connection between SNVs and hypoxia score. Only 4 genes were significantly different in the incidence of SNVs between the 2 groups (Figure 7b). Among them, the proportion of non-silent mutations only increased in ADAMTS19 in the high hypoxia score group. However, because the overall mutation frequency of these genes was not high, there was no statistically significant difference in the distribution of SNVs between the 2 groups. Therefore, we believe that hypoxia has a limited ability to induce SNVs. SNVs are essential for the development and progression of tumours. It is likely that important SNVs occur before the onset of hypoxia.

**Figure 7.**
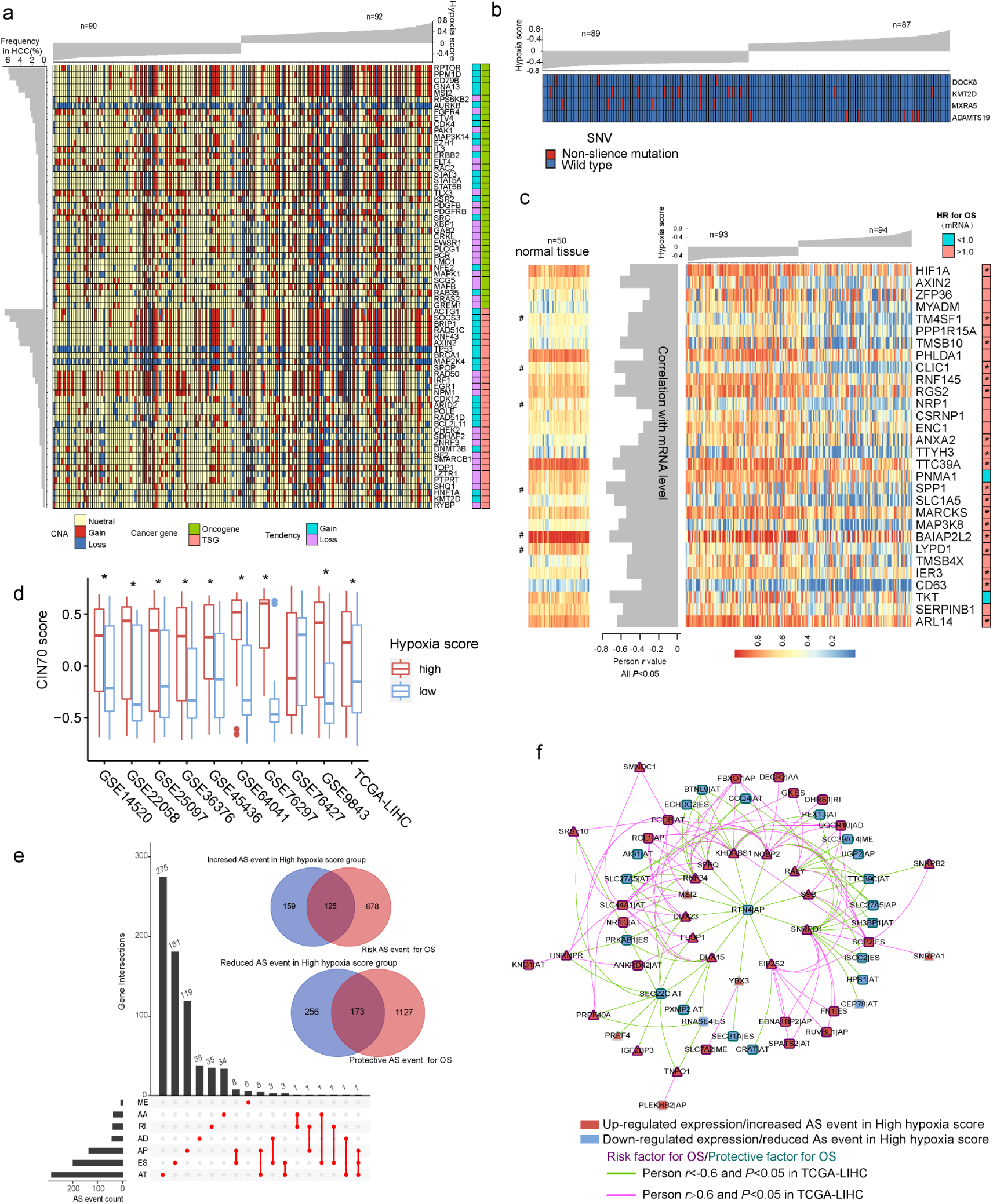
Differences in genomic and epigenetic alterations between groups with high hypoxia scores and low hypoxia scores. a) The difference in the incidence of copy-number aberrations (CNAs) in 71 cancer genes between the high hypoxia score group and the low hypoxia score group. b) The proportions of single-nucleotide variants (SNVs) in 4 genes are significantly different between the high hypoxia score group and the low hypoxia score group. c) Reductions in the methylation levels of 30 genes are accompanied by significant increases in the corresponding mRNA levels in the high hypoxia score group. The correlation between DNA methylation and corresponding mRNA expression was obtained through Pearson correlation analysis based on TCGA-LIHC data. The hazard ratios (HR) of the corresponding mRNAs for overall survival (OS) were calculated by the log-rank test for TCGA-LIHC data, and the cut-off was the median expression level. d) 70-gene chromosome instability (CIN70) was used to assess chromosome instability in tumour tissues from 10 hepatocellular carcinoma (HCC) datasets. The CIN70 scores are significantly different between tumour tissues with high hypoxia scores and low hypoxia scores. e) The occurrences of 713 AS events are significantly different between the high hypoxia score group and the low hypoxia score group. Some of the AS events are associated with the OS of HCC patients. f) The expression of 30 splicing factors in the high hypoxia score group and the low hypoxia score group are different, and their expression trends are consistent in 10 datasets. These splicing factors and the AS events with different occurrences between the 2 groups form a network. The correlations between the nodes of the network were calculated by Pearson correlation analysis based on TCGA-LIHC data. The relationship between the nodes and the OS of HCC patients was obtained through univariate Cox survival analysis.

**Figure 8.**
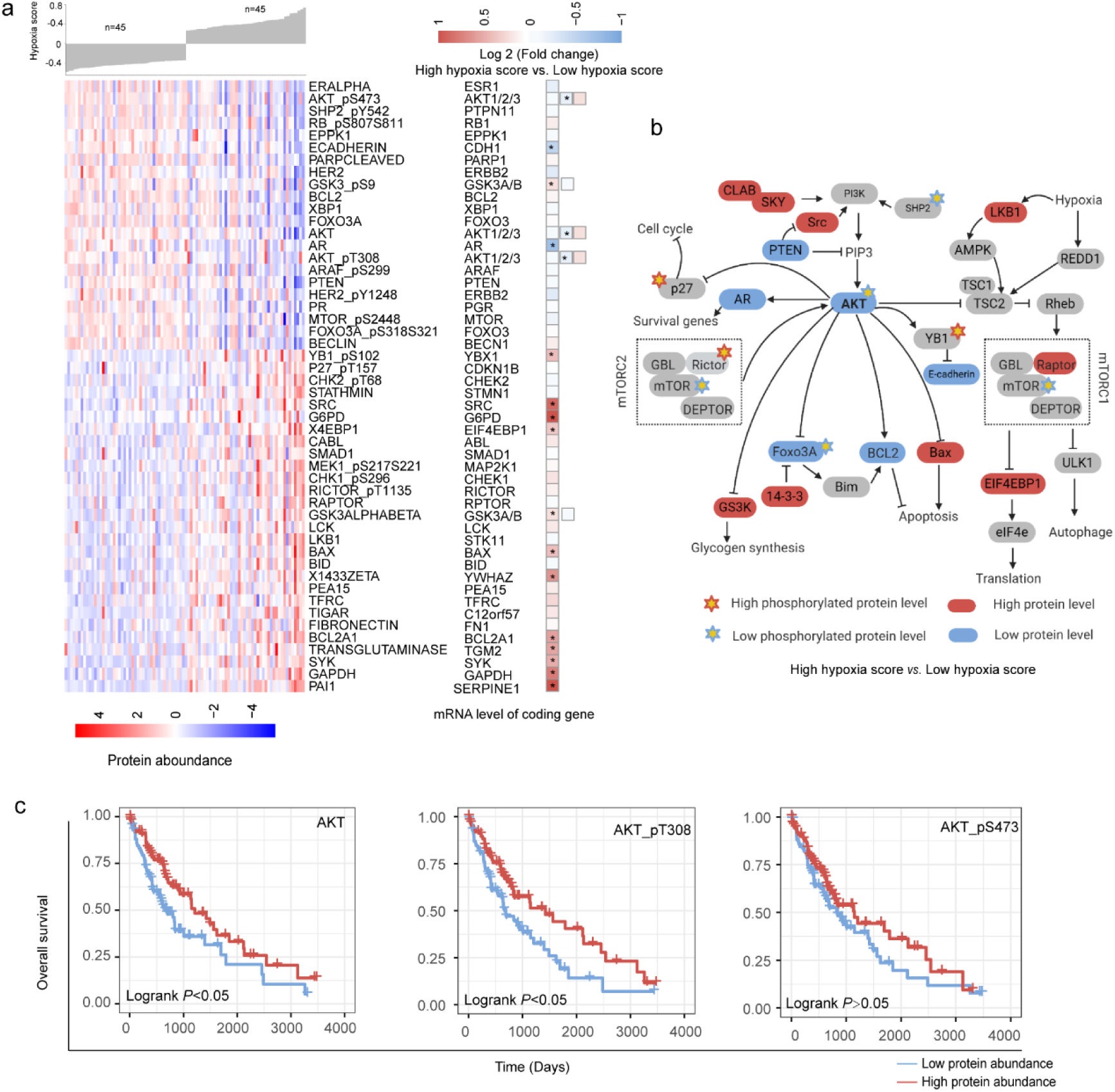
Changes in functional proteomics between hepatocellular carcinoma (HCC) patients with high hypoxia scores and low hypoxia scores. a) Fifty proteins with significant differences in abundance between HCC patients with high hypoxia scores and low hypoxia scores. b) A schematic diagram of some proteins with significant differences in abundance in the AKT/mTOR pathway. c The low abundance of AKT, AKT-pT308 and AKT_pS473 proteins in TCGA-LIHC indicates poor overall survival.

### Epigenetic alterations in HCC patients with different hypoxia scores

We found that there were significant differences in the methylation levels at 464 loci between patients with high hypoxia scores and patients with low hypoxia scores and that the methylation levels at most loci were significantly reduced in the high hypoxia score group. The methylation level increased at only a few loci in the high hypoxia score group. We jointly analysed methylation levels and mRNA expression levels and found a significant increase in mRNA expression levels of 30 genes in the high hypoxia score group (data from TCGA-LIHC) and a simultaneous decrease in their methylation levels (Figure 7c). The mRNA expression levels of these genes showed a significantly negative correlation with their methylation levels, and the high mRNA expression of most of these genes is a risk factor for HCC patient survival. Notably, the degree of methylation of HIF-1A was also reduced in the high hypoxia score group, which provides another explanation for the increase in HIF-1A mRNA expression in the high hypoxia score group. From the comparison of DNA methylation levels of the 30 genes in normal tissues and HCC tissues, the DNA methylation levels of some genes in normal tissues were not different from those in HCC tissues with low hypoxia scores but were higher than those in HCC tissues with high hypoxia scores (such as RGS2), and the DNA methylation levels of some other genes were highest in normal tissues, followed by those in HCC tissues with low hypoxia scores, and lowest in HCC tissues with high hypoxia scores (such as CLIC1). These results suggest that hypoxia exposure may promote the hypomethylation of some genes related to the onset and progression of HCC and that the hypomethylation of some other genes may be a characteristic feature of hypoxia exposure.

Chromosome instability occurs throughout the development and progression of tumours. The increase in chromosome instability will lead to HCC cell growth and enhanced invasiveness[35]. The 70-gene chromosome instability (CIN70) signature constructed by Carter et al. can effectively assess the chromosome instability of tumour cells[36]. We compared the CIN70 scores between patients with high hypoxia scores and patients with low hypoxia scores. The data showed that in 9 datasets, the CIN70 scores were significantly increased in the high hypoxia score groups (Figure 7d), suggesting that hypoxia exposure may cause chromosome instability in HCC cells.

Alternative splicing (AS) is an epigenetic feature that plays an important role in tumourigenesis and development[37]. Based on the TCGA-LIHC data, we found that the occurrences of 713 AS events (involving 681 mRNAs) differed significantly between the 2 groups and that most of these events were alternative terminator (AT), exon skipping (ES), and alternative promoter (AP) events (Figure 7e). In the high hypoxia score group, the percent spliced in (PSI) value of retained intron (RI) events of EPHB2 mRNA had the maximum reduction (adjusted *P* value < 0.001), and the AT events of GULP1 mRNA had the most increased PSI (adjusted *P* value < 0.01). The Cox analysis results suggested that the AS events with differences in occurrence between the 2 groups were related to HCC patient survival. The 681 mRNAs involved in differential AS events were subjected to functional enrichment analysis of biological processes and pathways. In addition to the HIF1 pathway and sugar and lipid metabolism pathways (indicating their relations to hypoxia exposure), these mRNAs are also involved in various classical pathways of tumourigenesis and development. In addition, we analysed the differences in expression of 404 splicing factors (SFs) between the high hypoxia score group and the low hypoxia score group. The differences in expression of 30 SFs were relatively consistent in 10 datasets (the condition of adjusted *P* < 0.05 was met in 5 or more datasets). Except for NRIP2, other SFs were upregulated in the high hypoxia score group. Next, correlations among these SFs and 713 AS events were analysed. When constructing the correlation network, we only set the thresholds at |Person r |> 0.6 and *P* < 0.05 and did not perform other node screening. Notably, in the obtained correlation network, each node reached logical harmony in variation trend and prognostic value (Figure 7f). For example, KHDRBS1 is an SF with elevated expression in the high hypoxia score group and is a risk factor for HCC patient survival. The occurrences of 7 AS events that were negatively correlated with KHDRBS1 were reduced in the high hypoxia score group, and these 7 AS events were protective factors for HCC patient survival. In contrast, 5 AS events that were positively correlated with KHDRBS1 increased in the high hypoxia score group, and these AS events were risk factors for HCC patient survival. Based on the evidence, we believe that AS events may be a response to hypoxia in HCC cells. Some hypoxia-induced changes in cell functions and behaviours may be achieved through the SF-AS network. The difference in AS between the high hypoxia score group and the low hypoxia score group may explain the prognosis difference between the 2 groups.

RNA N6-methyladenosine (m^6^A) modification is another important epigenetic feature of RNA[38]. The expression differences in 21 m^6^A regulators between the high hypoxia score group and the low hypoxia score group were analysed. In the 10 HCC datasets, most m^6^A regulators showed no significant difference between the 2 groups, but 1 m^6^A writer (WTAP), 1 m^6^A eraser (ALKBH5), and 1 m^6^A reader (YTHDF2) significantly increased in the high hypoxia score groups of 4 or more datasets.

### Changes in functional proteomics in HCC patients with different hypoxia scores

Based on the reverse phase protein array (RPPA) data of the TCGA-LIHC cohort, we analysed the differences in the abundance of 218 proteins between the high hypoxia score group and the low hypoxia score group. A total of 50 proteins exhibited differences in abundance between the 2 groups (Figure 8a). Increases in LKB1 and mTOR activity inhibition are hallmark events of hypoxia exposure, which once again confirmed the efficacy of our 21-gene signature in indicating hypoxia exposure. Approximately half of the protein changes were inconsistent with the changes in their mRNA levels, suggesting that the regulation of posttranslational levels by hypoxia cannot be ignored. The differentially expressed proteins were mainly concentrated in AKT/mTOR and its related pathways. The reduction in the AKT level and its phosphorylation level seems to be a core event. The change trends for some proteins were opposite to those reported by the relevant literature on AKT (Figure 8b), which suggests that in addition to the regulation of AKT, there may be a complex regulatory network that has not been discovered. Survival analysis (cut-off = median value, Figure 8c) showed that a low abundance of AKT, AKT-pT308 and AKT_pS473 indicated worse OS (except for AKT_pS473, log-rank *P* < 0.05). Hypoxia may induce some functions by inhibiting AKT in HCC. Overall, the changes in the proteins between the 2 groups reflected the dynamic balance of autophagy/apoptosis and the response to hypoxic stress to some extent. These proteins may affect the viability of the cells under hypoxia and their tolerance to hypoxia. Moreover, the altered protein abundance between the 2 groups may be related to the different drug sensitivities of HCC patients in the 2 groups. For example, the reduction in AKT_pS473 may cause the sensitivity of cells to sorafenib to decrease. Therefore, when formulating a treatment plan, it may be necessary to evaluate the hypoxic condition of the lesions.

### Different immunologic microenvironments in HCC patients with different hypoxia scores

The ESTIMATE algorithm was used to calculate the stromal and immune scores for patients in 11 HCC datasets. These 2 scores represent the infiltration of stromal cells and immune cells in tumour tissues. In GSE76297, GSE76427, and GSE9843, the stromal scores for tumour tissues were significantly higher in the high hypoxia score group than in the low hypoxia score group, whereas in other datasets, the stromal scores for tumour tissues were not significantly different between the 2 groups (Figure 9a). In GSE76297, GSE7642, GSE9843, GSE25097, GSE36376 and TCGA-LIHC, the immune scores for tissues with high hypoxia scores were significantly increased (Figure 9 b), and in other datasets (except for GSE10141), the immune scores for tissues with high hypoxia scores were also increased but not significantly. Almost all datasets showed that immune scores were higher in patients with high hypoxia scores than in patients with low hypoxia scores. The differences in immune scores between the 2 groups were significant in the GSE76297, GSE9843, GSE25097, GSE36376, and TCGA-LIHC datasets, suggesting that the degree of immune cell infiltration was higher in patients with high hypoxia scores than in patients with low hypoxia scores. We also used the CIBERSORT algorithm. The distribution of 22 types of immune infiltrating cells in HCC tissues in 11 datasets was evaluated. Figure 9c shows the different types of infiltrating immune cells in the high hypoxia score group and the low hypoxia score group. Here, we need to emphasize some results with high consistency in multiple datasets. In GSE14520, GSE22058, GSE25097, GSE36376, GSE9843, and TCGA-LIHC, a significant reduction in the proportion of infiltrating CD4 memory resting cells was accompanied by a significant increase in the proportion of CD4 memory activated cells in tissues with high hypoxia scores. In GSE14520, GSE22058, GSE25097, GSE45436 and TCGA-LIHC, a significant reduction in the proportion of infiltrating resting mast cells was accompanied by an increase in the proportion of activated mast cells in tissues with high hypoxia scores. Roughly speaking, the proportions of infiltrating M0 macrophages experienced different degrees of increase in tissues with high hypoxia scores in all datasets, but the differences were significant only in GSE14520, GSE22058, GSE25097, GSE36376 and TCGA-LIHC. In addition, the proportion of activated NK cells decreased in tissues with high hypoxia scores in almost all datasets but decreased significantly only in GSE10141, GSE14520 and GSE25097. These data suggest that hypoxia changes immune cell infiltration in the tumour tissues of HCC patients and that our 21-gene hypoxia signature can be used to distinguish the difference in infiltrating immune cells between tissues with different hypoxia scores. Finally, we analysed the expression of programmed death-1 (PD-1)/programmed death-ligand 1 (PD-L1) and cytotoxic T-lymphocyte antigen-4 (CTLA-4) in tissues with different hypoxia scores. In most datasets, there was no significant difference in PD-L1 levels between tissues with high hypoxia scores and tissues with low hypoxia scores. However, in GSE22052, GSE25097, GSE9843, and TCGA-LIHC, the expression of PD-L1 was consistently significantly higher in tissues with high hypoxia scores than in tissues with low hypoxia scores (Figure 9d). The ranks of PD-1 and CTLA-4 levels in tissues with high hypoxia scores and low hypoxia scores were different in different datasets (Figure 9e and f). Interestingly, in multiple datasets, the ranks of PD-1 levels in tissues with high hypoxia scores and low hypoxia scores were opposite to the ranks of CTLA-4 levels in tissues with high hypoxia scores and low hypoxia scores. For example, in GSE22058, PD-1 expression was higher in tissues with high hypoxia scores than in tissues with low hypoxia scores, but CTLA-4 expression was lower in tissues with high hypoxia scores than in tissues with low hypoxia scores. These data suggest that the 21-gene hypoxia signature may help to evaluate the response to immunotherapy in HCC patients.

**Figure 9.**
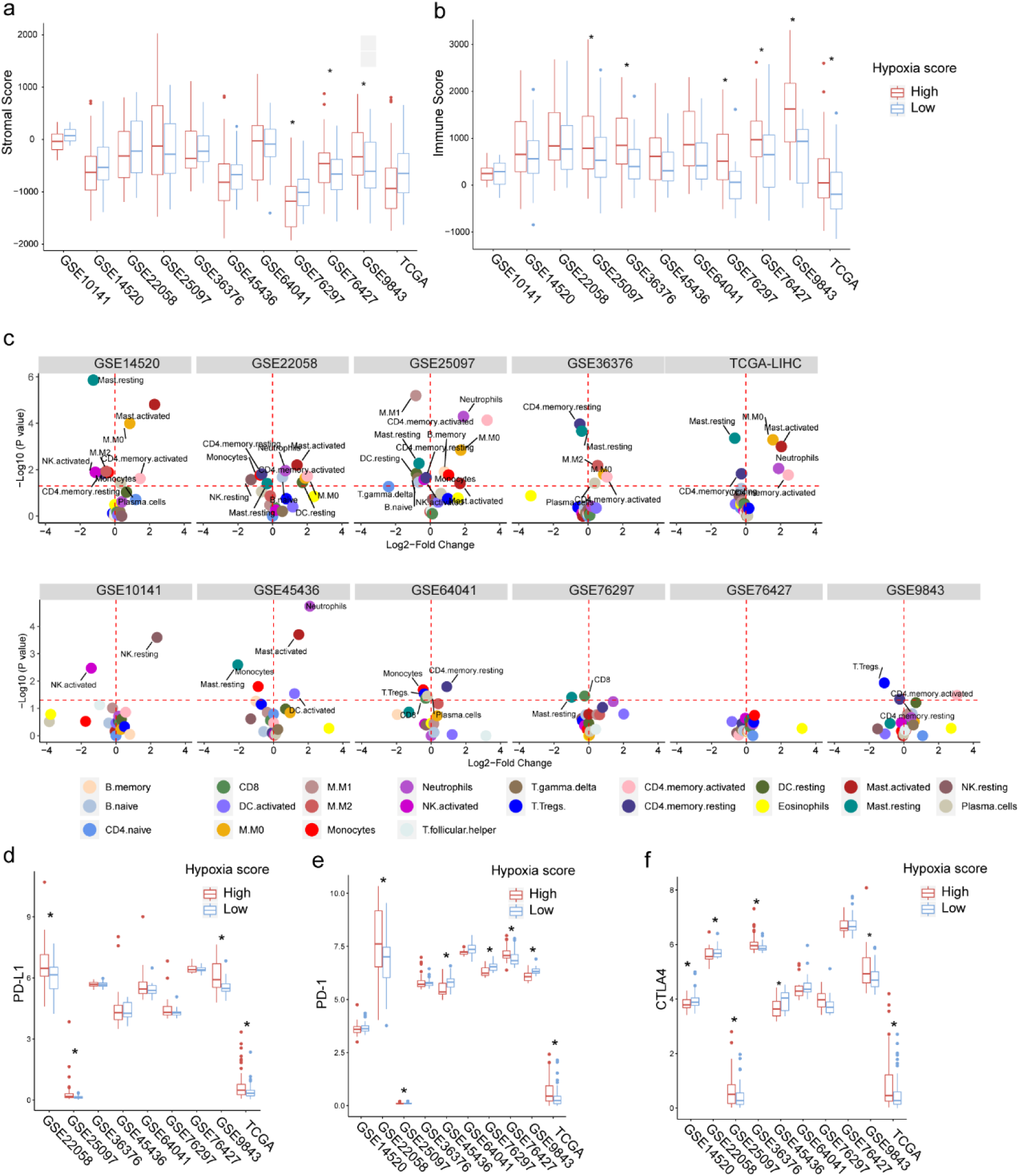
Differences in immune microenvironments between hepatocellular carcinoma (HCC) patients with high hypoxia scores and low hypoxia scores. a) and b) Differences in stromal scores and immune scores (obtained using the ESTIMATE algorithm based on mRNA data) for cancer tissues between HCC patients with high hypoxia scores and low hypoxia scores in 11 datasets. c) Differences in the types of infiltrating immune cells (obtained using the CIBERSORT algorithm based on mRNA data) in cancer tissues between HCC patients with high hypoxia scores and low hypoxia scores in 11 datasets. e) and f) Differences in the expression levels of programmed death-1 (PD-1), programmed death-ligand 1 (PD-L1) and cytotoxic T-lymphocyte antigen 4 (CTLA4) between HCC patients with high hypoxia scores and low hypoxia scores in 11 datasets. * compared with the low hypoxia score group, *P* < 0.05.

## Discussion

Hypoxia can change the gene expression level in tumour cells. Therefore, the detection of the levels of specific genes can indirectly reflect hypoxia exposure in tumour cells or tissues [39]. Although this method is not as accurate as the method using polarographic electrodes, it has high practicality. It can be predicted that with the development of high-throughput detection technology and improvements in related algorithms, the accuracy of hypoxia assessments based on gene signatures will improve, and hypoxia gene signatures have great potential in research and clinical application. In the past decade, different hypoxia gene signatures have been reported, and some excellent signatures, such as Buffas 15 -gene hypoxia signature, have proven to have the ability to indicate hypoxia in a variety of tumours [19]. Most previous studies only focused on the relationship between these signatures and the clinical characteristics and prognosis of cancer patients. Due to the explosive growth of high-throughput data and the development of the TCGA project, some recent studies have attempted to use these signatures to reveal molecular changes in tumour cells caused by hypoxia at multi-omic levels. For example, the pan-cancer study by Bhandari et al. showed molecular landmarks of tumour hypoxia across cancer types [14]. The integrative study by Ye et al. suggested that hypoxia-associated molecular features are closely related to the drugs used for tumour treatment [15]. The data obtained from these studies are instructive. However, none of the hypoxia gene expression signatures used in these studies were based on HCC, and the data on HCC used in these studies are also limited. Therefore, we believe that it is necessary to develop a new hypoxia gene signature that is HCC-specific and more robust than previous signatures. We believe that using this signature to specifically depict hypoxia-related molecular landscapes in HCC from a multi-omics perspective will benefit HCC patients.

In this study, we first used a microarray to obtain hypoxia-responsive mRNAs in 3 HCC cell lines and integrated them using the RRA algorithm to obtain a 21-gene hypoxia signature. Among the 21 genes, hypoxia-related genes CA9, HK2, and EGLN3 have been frequently reported [34, 40, 41]. Before further analysis, we performed cell-level validation in 3 other public datasets, demonstrating that our 21-gene hypoxia signature can indicate hypoxia exposure. At the tissue level, we found that our 21-gene hypoxia signature showed a good correlation with the 6 signatures reported (highly cited) by previous studies. Therefore, we believe that the novel 21-gene hypoxia signature has excellent robustness in the assessment of hypoxia exposure. After calculating hypoxia scores using the 21-gene hypoxia signature, we found that the hypoxia scores in the tumour tissues of HCC patients in 11 datasets were obviously grouped into 2 clusters, indicating that hypoxia exposure in HCC patients is not the same. In other words, some patients may respond to anti-hypoxia treatment while other patients do not need anti-hypoxia treatment because there is no excessive exposure to hypoxia. Currently, the efficacy of anti-hypoxia treatment is not satisfactory. Therefore, screening patients using a hypoxia signature before formulating a treatment programme may change this situation.

In recent years, the molecular classification of HCC has received attention. We used the 21-gene hypoxia signature to perform subtype analysis on HCC patients. In 2 independent cohorts (TCGA and GSE14520), the 21-gene hypoxia signature effectively classified patients into subtypes. The obtained subtypes differed regarding clinical characteristics, including stage and prognosis. Therefore, from the perspective of precision medicine, the classification of HCC based on hypoxia exposure has clinical implications for determining prognosis and developing personalized treatment plans, which may benefit specific patient groups. A model was constructed using hypoxia scores calculated based on the 21-gene hypoxia signature as an indicator. The model can well predict the OS rate of HCC patients and can indicate the recurrence of HCC. The hypoxia score was closely related to HCC stage, vascular invasion, and metastasis. These results were validated in multiple cohorts, suggesting the value of the 21-gene hypoxia signature in clinical application.

Next, we used the 21-gene hypoxia signature to describe hypoxia-related molecular landscapes in HCC tissues. The purposes of this study were to provide tissue-level evidence for explaining the mechanism of hypoxia in HCC, to provide a comprehensive overview of the role of hypoxia in HCC, and to find possible treatment or diagnostic targets. Considering the numerical distribution of hypoxia scores in tissues, we selected the upper quartile and the lower quartile to divide high hypoxia scores and low hypoxia scores. In this way, more differential changes may be obtained, creating a situation that is more in line with clinical practice. Because hypoxic conditions and molecular changes in tumour tissues are nonlinearly correlated, mild hypoxia exposure may not cause many molecular events. After comprehensive analysis, the patients in the high hypoxia score group and the patients in the low hypoxia score group were found to have many transcriptomic, genomic, epigenomic, and proteomic differences, including differences in mRNA, miR, and lncRNA expression, differences in CNAs, differences in DNA methylation levels, and differences in AS events. Some differential events are closely related to the prognosis of patients and are molecular mechanisms that explain the cancer-promoting effect of hypoxia. Therefore, these differential events can help screen HCC diagnostic and treatment targets. The results at multi-omic levels had some consistency, such as the simultaneous presence of high mRNA expression and decreased methylation of the corresponding DNA in the high hypoxia score group. However, some results were contradictory. For example, the GSEA results for the AKT/mTOR pathway based on mRNA level were not consistent with the proteomic results for the AKT/mTOR pathway. Therefore, it is necessary to investigate the molecular changes caused by hypoxia using multi-omics approaches. Based on the available data, we analysed the relationships between differential events and patient prognosis and extracted the useful data for translational medicine. In the supplemental files, we provide reference data with rich details, such as the lists of DE-mRNAs, DE-miRNAs, and DE-lncRNAs, the list of genes with CNAs, the list of genes with aberrant methylation loci, and the list of hypoxia-related RNA RA events. We hope that these data can inspire and help other researchers, improve research efficiency, and narrow the scope of research.

It should be emphasized that we found that the HIF-1A mRNA levels significantly increased in the high hypoxia score groups in the 10 datasets and showed a significantly positive correlation with hypoxia score. The reason why this point is emphasized is that most previous studies have focused on the posttranslational regulation of HIF-1A at the protein level, and some researchers even deliberately ignore changes in HIF-1A mRNA. Our data suggest that the regulation of HIF-1A mRNA levels cannot be ignored in hypoxia exposure, meaning that in treatment regimens targeting HIF-1A, not only HIF-1A protein but also HIF-1A mRNA should be targeted. In addition, our data showed that during hypoxia exposure, the level of HIF-1A might be regulated by a multilevel positive feedback network and that an mRNA/lncRNA network and methylation regulation might be part of this network.

In the last part of this study, the effect of hypoxia on immune cell infiltration patterns in HCC tissues was investigated. It was found that hypoxia induced the infiltration of immune cells in HCC tissues and that the proportion of specific types of immune cells changed. The current evidence is inadequate to clarify the role and significance of immune cells with changes in their proportions. However, based on existing reports, they might be involved in hypoxia-induced immune escape. Hypoxia may affect the expression levels of immune checkpoint regulators, such as PD-1 and PD-L1, suggesting that hypoxia may be related to the efficacy of immunotherapy.

The present study has some limitations. First, the use of a signature to reflect hypoxia exposure is indirect. Therefore, some differential events may not be directly caused by hypoxia. In addition to tumour cells, other cells are also present in HCC tissues. These confounding factors cannot be ruled out at present. In addition, the results need to be further validated by laboratory experiments. Finally, the number of genes in our signature should be further reduced for clinical application.

## Conclusions

In summary, the 21-gene signature developed in this study can effectively estimate hypoxia exposure in HCC tissues. The use of this signature can help to assess patient prognosis. Using this 21-gene signature, we described hypoxia-related molecular landscapes at the tissue level. The effect of hypoxia on HCC is complex, multilevel and networked. Different patients have different degrees of hypoxia exposure, and patients with different degrees of hypoxia exposure have different molecular characteristics and thus different responses to treatment. In the personalized treatment of HCC patients, the assessment of the degree of hypoxia is strongly recommended to benefit specific patient groups.

## List of Abbreviations

HCC: hepatocellular carcinoma
GEO: Gene Expression Omnibus
TCGA: The Cancer Genome Atlas
lncRNA: long noncoding RNAs
miR: microRNA
HIF: hypoxia inducible factor
RRA: robust rank aggregation
GSVA: gene set variation analysis
CAN: copy number alteration
SNV: single nucleotide variant
RPPA: The reverse phase protein array
RA: RNA alternative splicing
MCODE: Molecular Complex Detection
MSigDB: Molecular Signatures Database
GSEA: Gene set enrichment analysis
ESTIMATE: Estimation of Stromal and Immune cells in Malignant Tumor tissues

## Declarations

## Ethics approval and consent to participate

Not applicable.

## Consent for publication

Not applicable.

## Availability of supporting data

The datasets supporting the conclusions of this article are available in the TCGA data portal (https://portal.gdc.cancer.gov/) and the Gene Expression Omnibus (GEO, https://www.ncbi.nlm.nih.gov/geo/).

## Authors’ contributions

Study design: ZQN and LLP; Analysis and interpretation of data: ZQN, LJ, QLJ and LQ; drafting of the manuscript: ZQN, LPY, XMT and HXT; statistical analysis: LPY; data visualization: ZQN and LPY; funding and study supervision: LLP and LJ. All authors read and approved the final manuscript.

## Acknowledgements

Not applicable

## References

1. Bray F, Ferlay J, Soerjomataram I, Siegel RL, Torre LA, Jemal A: Global cancer statistics 2018: GLOBOCAN estimates of incidence and mortality worldwide for 36 cancers in 185 countries. CA Cancer J Clin. 2018; 68: 394–424.

2. Llovet JM, Burroughs A, Bruix J: Hepatocellular carcinoma. Lancet. 2003; 362: 1907–1917.

3. Kanwal F, Singal AG: Surveillance for Hepatocellular Carcinoma: Current Best Practice and Future Direction. Gastroenterology. 2019; 157: 54–64.

4. Fattovich G, Stroffolini T, Zagni I, Donato F: Hepatocellular carcinoma in cirrhosis: incidence and risk factors. Gastroenterology. 2004; 127: S35–50.

5. Llovet JM, Montal R, Sia D, Finn RS: Molecular therapies and precision medicine for hepatocellular carcinoma. Nat Rev Clin Oncol. 2018; 15: 599–616.

6. Lu C, Rong D, Zhang B, et al.: Current perspectives on the immunosuppressive tumor microenvironment in hepatocellular carcinoma: challenges and opportunities. Mol Cancer. 2019; 18: 130.

7. Jing X, Yang F, Shao C, et al.: Role of hypoxia in cancer therapy by regulating the tumor microenvironment. Mol Cancer. 2019; 18: 157.

8. Nobre AR, Entenberg D, Wang Y, Condeelis J, Aguirre-Ghiso JA: The Different Routes to Metastasis via Hypoxia-Regulated Programs. Trends Cell Biol. 2018; 28: 941–956.

9. Chen A, Sceneay J, Godde N, et al.: Intermittent hypoxia induces a metastatic phenotype in breast cancer. Oncogene. 2018; 37: 4214–4225.

10. Saxena K, Jolly MK: Acute vs. Chronic vs. Cyclic Hypoxia: Their Differential Dynamics, Molecular Mechanisms, and Effects on Tumor Progression. Biomolecules. 2019; 9.

11. Gilkes DM, Semenza GL, Wirtz D: Hypoxia and the extracellular matrix: drivers of tumour metastasis. Nat Rev Cancer. 2014; 14: 430–439.

12. Eltzschig HK, Carmeliet P: Hypoxia and inflammation. N Engl J Med. 2011; 364: 656–665.

13. van Malenstein H, Gevaert O, Libbrecht L, et al.: A seven-gene set associated with chronic hypoxia of prognostic importance in hepatocellular carcinoma. Clin Cancer Res. 2010; 16: 4278–4288.

14. Bhandari V, Hoey C, Liu LY, et al.: Molecular landmarks of tumor hypoxia across cancer types. Nat Genet. 2019; 51: 308–318.

15. Ye Y, Hu Q, Chen H, et al.: Characterization of Hypoxia-associated Molecular Features to Aid Hypoxia-Targeted Therapy. Nat Metab. 2019; 1: 431–444.

16. Haider S, McIntyre A, van Stiphout RG, et al.: Genomic alterations underlie a pan-cancer metabolic shift associated with tumour hypoxia. Genome Biol. 2016; 17: 140.

17. Kolde R, Laur S, Adler P, Vilo J: Robust rank aggregation for gene list integration and meta-analysis. Bioinformatics. 2012; 28: 573–580.

18. Hanzelmann S, Castelo R, Guinney J: GSVA: gene set variation analysis for microarray and RNA-seq data. BMC Bioinformatics. 2013; 14: 7.

19. Buffa FM, Harris AL, West CM, Miller CJ: Large meta-analysis of multiple cancers reveals a common, compact and highly prognostic hypoxia metagene. Br J Cancer. 2010; 102: 428–435.

20. Eustace A, Mani N, Span PN, et al.: A 26-gene hypoxia signature predicts benefit from hypoxia-modifying therapy in laryngeal cancer but not bladder cancer. Clin Cancer Res. 2013; 19: 4879–4888.

21. Ragnum HB, Vlatkovic L, Lie AK, et al.: The tumour hypoxia marker pimonidazole reflects a transcriptional programme associated with aggressive prostate cancer. Br J Cancer. 2015; 112: 382–390.

22. Sorensen BS, Toustrup K, Horsman MR, Overgaard J, Alsner J: Identifying pH independent hypoxia induced genes in human squamous cell carcinomas in vitro. Acta Oncol. 2010; 49: 895–905.

23. Winter SC, Buffa FM, Silva P, et al.: Relation of a hypoxia metagene derived from head and neck cancer to prognosis of multiple cancers. Cancer Res. 2007; 67: 3441–3449.

24. Bader GD, Hogue CW: An automated method for finding molecular complexes in large protein interaction networks. BMC Bioinformatics. 2003; 4: 2.

25. Yoshihara K, Shahmoradgoli M, Martinez E, et al.: Inferring tumour purity and stromal and immune cell admixture from expression data. Nat Commun. 2013; 4: 2612.

26. Roessler S, Jia HL, Budhu A, et al.: A unique metastasis gene signature enables prediction of tumor relapse in early-stage hepatocellular carcinoma patients. Cancer Res. 2010; 70: 10202–10212.

27. Camp RL, Dolled-Filhart M, Rimm DL: X-tile: a new bio-informatics tool for biomarker assessment and outcome-based cut-point optimization. Clin Cancer Res. 2004; 10: 7252–7259.

28. Chen P, Wang F, Feng J, et al.: Co-expression network analysis identified six hub genes in association with metastasis risk and prognosis in hepatocellular carcinoma. Oncotarget. 2017; 8: 48948–48958.

29. Hoadley KA, Yau C, Wolf DM, et al.: Multiplatform analysis of 12 cancer types reveals molecular classification within and across tissues of origin. Cell. 2014; 158: 929–944.

30. Dang K, Myers KA: The role of hypoxia-induced miR-210 in cancer progression. Int J Mol Sci. 2015; 16: 6353–6372.

31. Huang X, Le QT, Giaccia AJ: MiR-210--micromanager of the hypoxia pathway. Trends Mol Med. 2010; 16: 230–237.

32. Kishore S, Jaskiewicz L, Burger L, Hausser J, Khorshid M, Zavolan M: A quantitative analysis of CLIP methods for identifying binding sites of RNA-binding proteins. Nat Methods. 2011; 8: 559–564.

33. Xue Y, Ouyang K, Huang J, et al.: Direct conversion of fibroblasts to neurons by reprogramming PTB-regulated microRNA circuits. Cell. 2013; 152: 82–96.

34. Xu F, Yan JJ, Gan Y, et al.: miR-885-5p Negatively Regulates Warburg Effect by Silencing Hexokinase 2 in Liver Cancer. Mol Ther Nucleic Acids. 2019; 18: 308–319.

35. Eferl R, Trauner M: Chromosomal instability in HCC: a key function for checkpoint kinase 2. Gut. 2018; 67: 204–205.

36. Carter SL, Eklund AC, Kohane IS, Harris LN, Szallasi Z: A signature of chromosomal instability inferred from gene expression profiles predicts clinical outcome in multiple human cancers. Nat Genet. 2006; 38: 1043–1048.

37. Lu Y, Xu W, Ji J, et al.: Alternative splicing of the cell fate determinant Numb in hepatocellular carcinoma. Hepatology. 2015; 62: 1122–1131.

38. Chen Y, Peng C, Chen J, et al.: WTAP facilitates progression of hepatocellular carcinoma via m6A-HuR-dependent epigenetic silencing of ETS1. Mol Cancer. 2019; 18: 127.

39. Yang L, Roberts D, Takhar M, et al.: Development and Validation of a 28-gene Hypoxia-related Prognostic Signature for Localized Prostate Cancer. EBioMedicine. 2018; 31: 182–189.

40. Yu SJ, Yoon JH, Lee JH, et al.: Inhibition of hypoxia-inducible carbonic anhydrase-IX enhances hexokinase II inhibitor-induced hepatocellular carcinoma cell apoptosis. Acta Pharmacol Sin. 2011; 32: 912–920.

41. Li S, Rodriguez J, Li W, et al.: EglN3 hydroxylase stabilizes BIM-EL linking VHL type 2C mutations to pheochromocytoma pathogenesis and chemotherapy resistance. Proc Natl Acad Sci U S A. 2019; 116: 16997–17006.

